# Altered neuronal start codon stringency favors cap-independent repeat-associated non-AUG translation

**DOI:** 10.64898/2026.07.16.739042

**Authors:** Clare M. Wieland, Shannon E. Wright, Sydney Willey, Ishita Purwar, Samantha J. Grudzien, Amy Krans, Erinn Laimon, Melissa J. Asher, Adrian Isaacs, Amanda L. Garner, Peter K. Todd

## Abstract

Intronic GGGGCC repeat expansions in *C9orf72* cause amyotrophic lateral sclerosis (ALS) and frontotemporal dementia (FTD). This expansion supports a non-canonical form of translational initiation known as repeat-associated non-AUG (RAN) translation to produce toxic dipeptide repeat proteins that contribute to neurodegeneration. Here, we find that the efficiency of RAN translation and its dependency on the 5′ 7-methylguanosine mRNA cap are variable across cell types, with both rodent neurons and human iNeurons favoring cap-independent RAN translation from two distinct repeats (CGG and GGGGCC) across multiple reading frames. Treatment with an eIF4E inhibitor that blocks global cap-dependent translation enhances RAN translation specifically in neurons. Intriguingly, cap-independent RAN translation exhibits less reliance on near-cognate codons for initiation than cap-dependent RAN translation. This finding led us to identify a surprising global increase in start codon stringency in neurons as a contributor to the relatively higher cap-independent RAN translation in this cell type. This effect correlates with a cytoplasmic redistribution of eIF1 in neurons and is reversed with neuronal overexpression of the eukaryotic initiation factor eIF5, which relaxes start codon stringency and selectively enhances cap-dependent RAN translation. Taken together, these findings reveal several neuron-specific features of translational regulation that favor cap-independent RAN translation with implications for nucleotide repeat expansion disorder pathogenesis and neuronal translational regulation.

**GRAPHICAL ABSTRACT:** 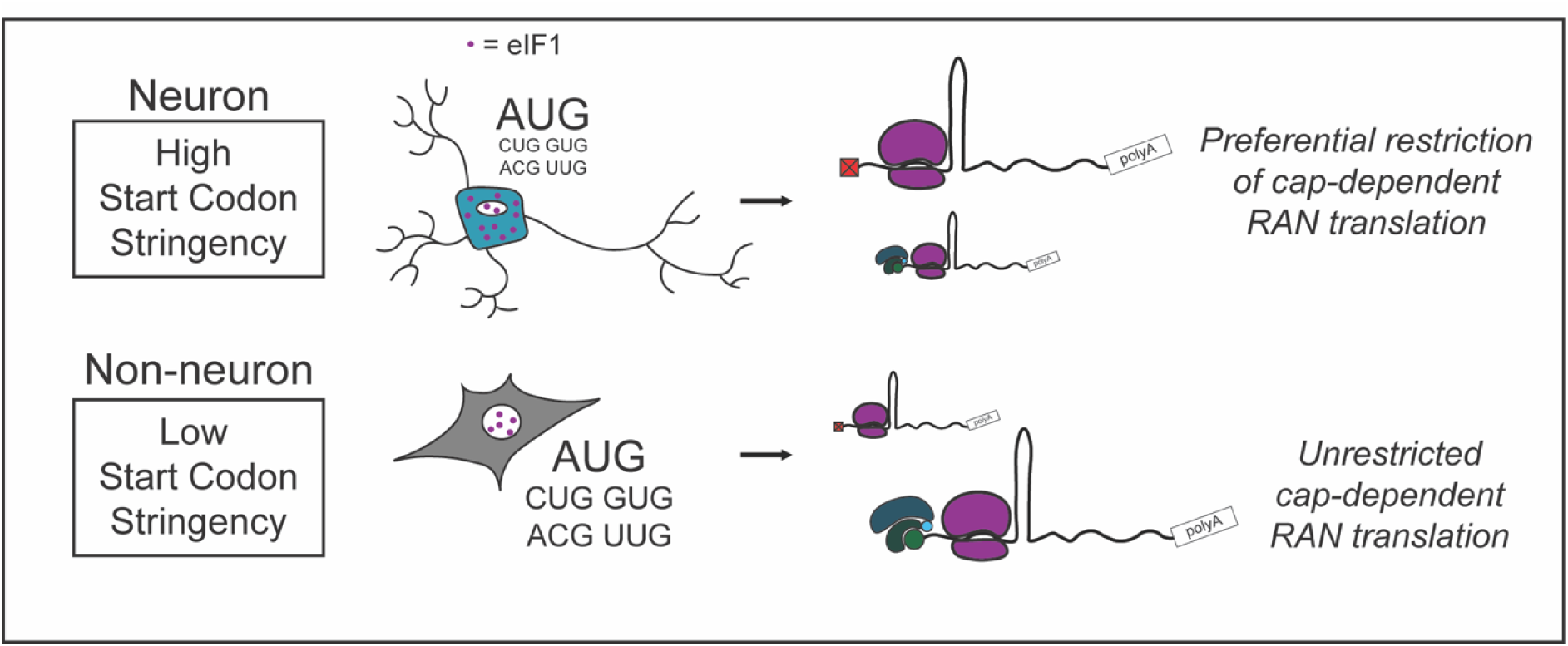

## INTRODUCTION

Short tandem repeat expansions contribute to over sixty predominantly neurologic disorders (1–3), including Huntington’s disease, Fragile-X associated Tremor/Ataxia Syndrome (FXTAS), and *C9orf72* associated amyotrophic lateral sclerosis (C9-ALS) and frontotemporal dementia (C9-FTD) (2). In C9- ALS/FTD, a hexanucleotide GGGGCC repeat in the first intron of *C9orf72* drives pathology. Healthy individuals typically have less than 30 repeats while patients typically have several hundred or even thousands of repeats, although disease has been observed in patients with as few as thirty repeats (1, 4). Repeat expansions in *C9orf72* are the most common known cause of both ALS and FTD (5–7).

Repeat expansions, including C9-ALS/FTD, support a noncanonical mode of translational initiation known as repeat-associated non-AUG (RAN) translation (1, 2, 8). Canonical translation initiation is a highly regulated process (9–11). Translation begins when the cap-binding complex eIF4F (including eIF4E, which directly binds the cap) and polyadenylate-binding protein (PABP) bind to the 5′ m^7^G-cap and the polyadenylated tail of mature mRNA, respectively. Once activated, the mRNA is recruited to the 43S translation pre-initiation complex (PIC, which contains the 40S ribosomal subunit, several initiation factors, and methionine initiator tRNA). The PIC binds to eIF4F and the activated mRNA, forming the 48S translation initiation complex. This complex then scans the mRNA in a 5′-3′ direction until an AUG start codon in an appropriate sequence context (known as the “Kozak context”) is reached (9–11). Start codon recognition releases most initiation factors and allows for recruitment of the 60S ribosomal subunit, formation of the 80S initiation complex, and the transition to elongation with the first peptide bond (9, 12, 13). Fidelity of start codon recognition is governed by the dynamic ratio of eIF1, which enhances start codon stringency, and eIF5, which loosens it, enhancing non-canonical forms of translation including RAN translation (9, 14, 15). Other initiation factors; including eIF1A, eIF5B, eIF5MP and eIF2; play additional roles in governing start codon selection (9). Secondary structure in the 5’ UTR also has profound effects on start codon selection. Secondary structure upstream of the target AUG start codon inhibits start codon selection through steric hindrance of cap binding or by impeding scanning (16–18). Downstream secondary structure increases initiation including at non-cognate codons or those in poor Kozak context (19–24). In concert, mRNA structure and initiation factor activity tightly regulate start codon selectivity, with profound effects on protein expression.

RAN translation uses two non-exclusive mechanisms. First, translation can occur via a scanning mechanism that shares many features with canonical translation: a 5′ m^7^G-cap is required, and scanning occurs until the 48S complex reaches the secondary structure formed by the often GC-rich repeat. There, the 48S complex stalls and can use a near-cognate (almost AUG, such as CUG or ACG) codon to recruit the 60S ribosomal subunit and begin translation (22, 25–28). Alternatively, RAN translation can occur in a cap-independent manner through a poorly understood process that is thought to utilize an internal ribosome entry site (IRES) (29–35). In viral IRES-mediated translation, complex RNA secondary structures and IRES trans-activating factors (ITAFs) can bypass the need for a 5′ cap, some canonical initiation factors, and an AUG codon and allow for recruitment of the ribosome with or without scanning along the mRNA (36, 37). The mechanism by which cap-independent RAN translation occurs is unknown but may involve specific extension segment ribosome protein factors such as RPS25 (33).

C9-ALS/FTD associated RAN translation (C9-RAN) produces dipeptide repeat proteins (DPRs) that accumulate in patient brain and spinal cord (38–42) and are neurotoxic (43–51). C9-RAN occurs in all three reading frames on both the sense and antisense strands, producing 5 distinct DPRs: poly-glycine-alanine (GA), poly-glycine-proline (GP), and poly-glycine-arginine (GR) from the sense strand; and poly-proline-arginine (PR), poly-proline-alanine (PA), and a second GP from the antisense strand (8, 38, 39, 41, 52). Cellular stress enhances DPR production, leading to a feed forward loop that is thought to contribute to disease (25, 31, 32, 53). Although the exact mechanisms of C9-RAN are still unclear, numerous groups have shown that the GA DPR is the most abundant DPR in patient brains and the most efficiently generated DPR in model systems (25, 26, 31, 32, 53, 54). RAN translation of GA DPRs is highly dependent on a near cognate CUG codon located 24 nucleotides upstream of the repeat in a strong Kozak context, which is in frame for GA but not GP or GR (14, 25, 26, 32).

In reporter assays and simple cell-based systems, C9-RAN translation is largely a cap-dependent and eIF4A-scanning dependent initiation process (25–28). However, most of these previous studies utilized either in-vitro cellular lysate systems, such as rabbit reticulocyte lysates, or human immortalized cell lines (e.g., HEK-293T cells). Here, we re-examine C9-RAN translation and uncover a surprisingly high degree of cap-independent translation in both rodent neurons and human iPSC-derived neurons (iNeurons). In neurons, inhibition of the cap-binding factor eIF4E enhances endogenous C9-RAN translation while inhibiting global translation. This observed shift towards cap-independent C9-RAN translation is accompanied by a higher degree of start codon stringency in neurons which supports a greater proportion of cap-independent RAN translation in this cell type.

Together, these findings reveal several features of translational regulation specific to neurons that favor cap-independent RAN translation with implications for broader nucleotide repeat expansion disorder pathogenesis.

## MATERIAL AND METHODS

### RNA synthesis

RNA was in vitro transcribed from linearized plasmid reporters. Firefly reporters and eIF5 were linearized using PmeI (NEB R0560); all other reporters were linearized using PspOMI (NEB R0653). RNA was transcribed using the HiScribe® T7 High Yield RNA Synthesis Kit (NEB E2040), with either functional, ARCA-cap (3′-O-Me-m^7^G(5’)ppp(5’)G RNA Cap Structure Analog, NEB S1411) or nonfunctional, A-cap (m^7^G(5’)ppp(5’)A RNA Cap Structure Analog, NEB S1405) which stabilizes against degradation but does not allow for cap binding. T7 reactions were run at 37°C for 2 hours. Reactions were treated with 2 units DNAse I (RNAse free) (NEB M0303) to degrade the remaining DNA template. RNA was polyadenylated using the *E. coli* Poly(A) Polymerase kit (M0276), with the reaction running at 37°C for 1 hour. RNA was then clean and concentrated using the RNA Clean & Concentrator-25 kit (Zymo Research 1018). RNA size and quality was assessed on denaturing agarose gel.

### Cell Culture

*HEK293* cells were ordered from American Type Culture Collection (ATCC) and cultured in DMEM with high glucose (Gibco 11965118), supplemented with 10% fetal bovine serum (FBS, Bio-Techne S11150) and 1% penicillin-streptomycin (Gibco 15070063). Cells were maintained at 37°C, 5% CO2.

*HeLa* cells were ordered from American Type Culture Collection (ATCC) and cultured in DMEM with high glucose (Gibco 11965118), supplemented with 10% fetal bovine serum (FBS, Bio-Techne S11150), 1% penicillin-streptomycin (Gibco 15070063), and 1% NEAA non-essential amino acids (Gibco 11140050). Cells were maintained at 37°C, 5% CO2.

*Rat primary hippocampal neurons* were dissected from P0-P1 Sprague-Dawley rat pups of both sexes. Hippocampi were isolated and dissociated using papain as in Sutton et al., 2006 (55). Cells were plated in neuron growth media (NGM): Neurobasal-A (Gibco 10888022), 1x GlutaMax (Gibco 35050061), and 1x B27 Supplement (Gibco 17504044) and maintained at 37°C, 5% CO2.

*iPSCs* were obtained from American Type Culture Collection (ATCC) as fibroblasts (ATCC PCS-201- 010) and differentiated into iPSCs as in Maltby et al., 2024 (56). Poly-GA HiBiT iPSC line was obtained from Adrian Isaac’s lab at UCL. Cells were cultured in mTeSR plus (Stem Cell, 100-0276) or TeSR-E8 (Stemcell 05990) and maintained at 37°C, 5% CO2.

*iNeurons* were differentiated as in Miller et al., 2025 (28). Briefly, a neurogenin-2 (Ngn2) overexpression cassette was delivered into iPSCs. iPSCs were cultured in TeSR-E8 (Stemcell 05990) media plus 2μg/mL doxycycline for 2 days and frozen for further use. To begin iNeuron differentiation (day 1), cells were thawed onto dPGA/Geltrex coated plates in TeSR-E8 containing doxycycline and Rock inhibitor. On day two, media was changed to transition media containing N2 with 1x NEAA supplement (Gibco 11140050), 10 ng/mL BDNF, 10 ng/mL NT3, 0.2 μg/mL laminin, 2 μg/mL doxycycline in half TeSR-E8 media, half DMEM F12 (Gibco 11320033). From day 3 until maturity (day 14), iNeurons were cultured in maintenance media which contained 1x B27 supplement (Gibco 17504044), 1x Glutamax supplement (Gibco 35050061), 10 ng/mL BDNF, 10 ng/mL NT3, 0.2 μg/mL laminin, and 1x Culture One in Neurobasal-A (Gibco 10888022). Cells were maintained at 37°C, 5% CO2 and underwent a 50% media change twice weekly.

*iAstrocytes* were differentiated as in Tcw et al., 2017 (57). Briefly, iPSCs were differentiated into neural progenitor cells (NPCs) via dual-SMAD inhibition protocol as in Maltby et al., 2024 (56). Low passage NPCs were then differentiated in astrocyte medium (ScienCell 1801) for 30 days with media changes every 48 hours. After differentiation, astrocytes were maintained in astrocyte medium at 37°C, 5% CO2.

### Immunocytochemistry

iNeurons were differentiated, as above, on dPGA/Geltrex-coated glass chamber slides until day fourteen. iAstrocytes were plated onto dPGA/Geltrex-coated glass chamber slides. The next day, cells were washed with PBS containing magnesium and calcium three times and fixed with 4% PFA for 15 minutes. Cells were washed again and permeabilized with 0.1% Triton X-100 for 5 minutes before blocking with 5% normal goat serum for 1 hour. Cells were then incubated with the following primary antibodies at 4°C overnight: GFAP at 1:500 dilution (75-240, NeuroMab), MAP2 at 1:1000 dilution (AB5622, Millipore), eIF1 at 1:500 dilution (12496S, CST), and eIF5 at 1:250 dilution (13894S, CST). For co-staining with eIF1 and eIF5 in neurons, MAP2 at 1:500 dilution (M1406, Sigma) was used. Cells were washed again and incubated for 1 hour at room temperature in the dark with Alexa Fluor secondary antibodies at 1:500 dilution (Fisher). Cells were washed again before staining and mounting with a coverslip using ProLong Gold with DAPI (Fisher). Images were captured at the same exposure for each cell line at 20x magnification with a Leica Stellaris 5 and LAS X software. Brightness and contrast were adjusted linearly to improve visualization in the representative images.

### Image Quantification

Imaging of ICC was performed at 20x magnification with a Leica Stellaris 5 and LAS X software. Channels were imaged sequentially to minimize bleed-through. Cells were imaged in a series of 15-25 1μm Z-planes to resolve the entire nucleus. Images were analyzed using ImageJ. Quantification of nuclear and cytoplasmic intensity was performed on maximum projections of scans. Intensity thresholding of the DAPI signal was used to identify nuclei ROIs. The resulting ROIs were applied to the image and the signal outside the nucleus was cleared before measuring average nuclear intensity across all cell nuclei in the image. To measure the cytoplasmic signal, the average intensity was measured across the entire cell for all cells in the image. The resulting nuclear signal was divided by the cytoplasmic signal for a given image to calculate its nuclear/cytoplasmic ratio. This was repeated for 5 images total for each cell line from two independent stainings. Data were analyzed by unpaired t-test with Welch’s correction.

### RNA Transfections

For nanoluciferase assay, cells were plated and transfected as follows. HEK293 cells were plated at 2x10^4^ cells per well in a 96 well plate in maintenance media without antibiotic. HeLa cells were plated at 1x10^4^ cells per well in a 96 well plate in maintenance media without antibiotic. iAstrocytes were plated at 0.5x10^4^ cells per well in a 96 well plate in maintenance media. Transfection was performed as below 24 hours after plating. iNeurons were differentiated in 96 well plates, as above, with 0.75x10^4^ cells per well and transfected on day fourteen. Rat primary hippocampal neurons were plated at 3.5x10^4^ cells per well in a 96 well plate on day of dissection and transfected 7-8 days later.

Cells were transfected with 50 ng nanoluciferase reporter and 50 ng firefly reporter using Lipofectamine MessengerMAX reagent according to manufacturer protocol (Thermo Fisher, LMRNA008). For eIF5 overexpression experiments, 100 ng of eIF5 RNA was cotransfected with 50 ng nanoluciferase reporter and 50 ng firefly reporter. After 24 hours, nanoluciferase assay was performed, as below.

For Western blot analysis, cells were plated and transfected as follows. HEK293 cells were plated at 1x10^5^ cells per well in a 12 well plate in maintenance media without antibiotic. HeLa cells were plated at 0.5x10^5^ cells per well in a 12 well plate in maintenance media without antibiotic. iAstrocytes were plated at 0.5x10^5^ cells per well in a 12 well plate in maintenance media. Transfection was performed as below 24 hours after plating. iNeurons were differentiated as above in 12 well plates with 250,000- 500,000 cells per well and transfected on day 13-14. Rat primary hippocampal neurons were plated at 3.5x10^5^ cells per well in a 12 well plate on day of dissection and transfected 7-8 days later.

Cells were transfected with 500 ng nanoluciferase reporter using Lipofectamine MessengerMAX reagent according to manufacturer protocol (Thermo Fisher, LMRNA008). After 24 hours, cells were lysed for western blot analysis, as below.

### Lentivirus

A lentiviral overexpression plasmid was purchased from Vector Builder. GFP control vector (pLLEV- GFP) was purchased from the University of Michigan Vector Core. Lentiviruses were packed at the University of Michigan Vector Core with HIV lentiviruses and then concentrated to 10x concentration in 10 mL of DMEM. To transduce iNeurons, media was removed from iNeurons on DIV13, and a half media change with diluted control GFP lentivirus or eIF5 overexpression lentivirus in maintenance media was applied. Twice weekly media changes were performed, as above. For Western blotting, cells were harvested on DIV23, as below, to confirm eIF5 overexpression. For HiBit assay, cells were analyzed on DIV23, as below. For RNA transfections, RNA was transfected, as above, on DIV22 and nanoluciferase assay was performed on DIV23, as above.

### Nanoluciferase Assay

Cells were lysed in 70 uL Glolysis Buffer (Promega E26661) for 5 minutes at room temperature on a shaker. NanoGlo Substrate was mixed in a 1:50 ratio with NanoGlo Buffer (Promega N1150).

Subsequently, cell lysate was moved to opaque 96 well plates (black for HEK293 and HeLa cells, white for all others) and was shaken in the dark at room temperature for 5 minutes in a 1:1 volumetric ratio with the mixed NanoGlo Solution to assess for nanoluciferase signal, One-Glo Solution (Promega E6130) to assess for firefly signal, or CellTiter-Glo Solution (Promega G7570) to assess for cell viability. A GloMax 96 Microplate Luminometer was used to quantify luminescence.

### HiBiT Assay

Poly-GA HiBiT iNeurons were differentiated as above with a density of 30,000 cells/well in a 96 well plate and allowed to mature for 14 days. On day fourteen, HiBiT detection reagent was prepared according to manufacturer instructions by diluting LgBiT Protein 1:100 and Nano-Glo HiBiT Lytic Substrate 1:50 in Nano-Glo HiBiT Lytic Buffer (Promega N3030). Freshly prepared reagent was added directly to the cells in a 1:1 ratio with culture media and incubated for 10 minutes at room temperature in the dark on a shaker. Cell viability was determined by removing media and performing cell titer assay as above. After incubation, cell lysate mix was transferred to black opaque 96 well plates and luminescence was quantified using a GloMax 96 Microplate Luminometer.

### Western Blotting

Cell lysis was performed as follows. Media was removed and cells were washed once in 300 uL of cold PBS. Cells were lysed in 120-150 uL of cold RIPA buffer with Complete Mini protease inhibitor tablet (Roche, 11836153001) for 30 minutes at 4°C. Lysate was mixed with 6x sample loading dye and 25:3 2-mercaptoethanol, boiled at 95°C for 5 minutes, and homogenized with a Series 60 Sonic Dismembrator (Thermo Fisher) for 25 seconds. Samples were stored at -20°C before use. The SUnSET assay was performed as in Schmidt et al., 2009 (58).

Samples were run on Novex Tris-Glycine Mini Protein Gels, 12%, 1.0 mm (ThermoFisher XP00125) or 4-20%, 1.0 mm (Thermofisher XP04205) with the following antibodies and conditions: 1:1000 ANTI-FLAG M2 antibody, Mouse monoclonal (mouse, Sigma F1804), 1:500 alpha Tubulin Monoclonal Antibody (YL1/2) (rat, ThermoFisher MA1-80017), 1:500 Anti-Puromycin Antibody, clone 12D10 (mouse, Sigma MABE343), 1:1000 GAPDH (Proteintech 60004-1-Ig), 1:1000 Actin (Sigma A1978), 1:1000 ATF-4 (SCBT sc-200), 1:1000 eIF1 (CST 12496S), 1:1000 eIF5 (D5G9) (CST 13894S), 1:1000 RPS11 (abcam ab157101), overnight at 4°C on a rocker. HRP goat anti-mouse (Jackson 115-035-146) or goat anti-rat (Jackson 112-035-003) secondary antibody was applied at 1:5000 or 1:10000 for 1 hour at room temperature on a rocker and bands were visualized on film. For loading control (alpha tubulin), Alexa Flour IR rat 800 (LiCOR 926-32219) secondary was used at 1:5000 for 1 hour at room temperature and bands were visualized on the Odyssey DLx (LICORbio). Band intensities were measured using ImageJ and normalized to alpha-tubulin.

### Drug Treatments

For inhibition of eIF4E, eFT-565 (565) (59, 60) was generously provided by Amanda L. Garner, PhD. Cells were treated with 0-15 μM eFT-565 dissolved in DMSO. For transfection-based assays, cells were plated as above, treated with the drug or vehicle, incubated for 4.5 hours at 37°C, 5% CO2, transfected, and incubated for another 19.5 hours before the nanoluciferase assay, as above.

For HiBiT assays, cells were treated with 7.5 μM eFT-565, 10 μg/mL puromycin (Sigma P8833), or vehicle and incubated for 24 hours at 37°C prior to assay, as above. The SUnSET assay was performed as in Schmidt et al., 2009 (58), with 24 hours of 7.5 μM eFT-565 treatment, 24 hours of vehicle treatment, or 15 minutes of 100μg/mL cycloheximide (Sigma 01810) treatment. Effect on translation was determined via western blot, as above.

For activation of the integrated stress response, cells were treated with either 2μM thapsigargin (Thermo T7459) dissolved in DMSO, 20 μM sodium arsenite (Fluka 35000) dissolved in water, or vehicle. 1 hour later, transfection was performed as above, and 5 hours later nanoluciferase assay was performed as above. For confirmation of the activation of the integrated stress response, cells were treated with either 2μM thapsigargin (Thermo T7459) dissolved in DMSO, 20 μM sodium arsenite (Fluka 35000) dissolved in water, or vehicle and 6 hours later lysed for Western blot, as above.

### Allen Brain Atlas RNA-seq Data Acquisition

Single-nucleus RNA sequencing datasets were obtained from Allen Brain Atlas RNA-seq Data Portal, provided by the Allen Institute for Brain Science. Data came from Human M1 10x dataset and Human Multiple Cortical Areas SMART-seq downloaded on October 23, 2025.

The Human M1 10x dataset consists of 76,533 nuclei derived from two post-mortem human primary motor cortex. The Allen Institute quantified gene expressions that consist of both intronic and extronic reads. The Human Multiple Cortical Areas SMART-seq dataset consists of 49,494 single nuclei isolated from middle temporal gyrus, anterior cingulate gyrus, primary visual cortex, primary somatosensory cortex, and primary auditory cortex. The Allen Institute quantified aggregated gene-level exon and intron counts.

For both datasets, Allen Institute-provided trimmed-means.csv files were used. These files contain transcriptomic clusters as columns and genes as rows with trimmed means as values. The Allen institute calculated trimmed-means by first normalizing gene expression as the log2 counts per million of the total exonic and intronic counts. Expression values in the upper and lower 25% of values were excluded, and the average of the remaining middle 50% values were averaged to get trimmed means.

In accordance with the Allen Institute citation policy, the Human M1 10x dataset was cited together with Bakken et al. (2021), and the Human Multiple Cortical Areas SMART-seq dataset was cited together with Tasic et al. (2018) and Hodge et al. (2019).

### RNA-seq Data Processing and Analysis in R

R-analyses were performed using the tidyverse, dyplr, and tidyr packages. The trimmed-mean matrices were imported into R, and rows with zero transcriptomic clusters were removed. Genes of interest were defined as EIF1, EIF5, BZW2, RPS11, EIF5B, EIF1AX, and ACTB. To be consistent with Allen Institute, their gene symbols were retained in the output files. Data analyses of EIF1A correspond to EIF1AX, and EIF5MP corresponds to BZW2.

For each dataset, we extracted the transcriptomic cluster and transformed the data so that the transcriptomic clusters were represented in rows. Broad cell classes were defined based on the characters included in the Allen Institute cluster labels: “Inh” = “Inhibitory”, “Exc” = Excitatory, “Astro” = Astrocytes, “Oligo” = Oligodendrocytes, and “Micro” = Microglia. Clusters that did contain these labels were excluded, and further analysis focused on astrocytes, excitatory neurons, and inhibitory neurons.

For each transcriptomic cluster, expression ratios of the genes of interest were calculated relative to ACTB expression. Ratio data from both datasets were combined into an individual table for each separate ratio. These ratio tables were exported as comma separate valued for downstream visualization and analysis.

For transcriptomic analyses, Kolmogorov-Smirnov test was used to assess distribution differences between groups. Graphs represent mean +/- SEM. N = 226 Neurons, N = 6 Astrocytes.

### Biological Resources

*Cell Lines*: HEK293 (ATCC CRL-1573), HeLa (ATCC CCL-2), Fibroblasts (ATCC PCS-201-010), poly-GA HiBiT iPSC line (Adrian Isaacs Lab, UCL)

*Organisms*: Sprague-Dawley Rats (Charles River, 001)

### Statistical Analyses

Statistical analyses were performed using GraphPad Prism, with specific analyses indicated in figure legends. Briefly, for experiments with two conditions a two-tailed Student’s t test with Welch’s correction was performed. For experiments with more than two conditions, a Brown-Forsythe one-way ANOVA with Dunnet’s multiple comparison correction was performed. For transcriptomic analyses, Kolmogorov-Smirnov test was used to assess distribution differences between groups.

Graphs represent mean ± SEM. N for each experiment is listed in the figure legends and represents biological replicates. All data presented represents results from at least three independent experiments.

### Reagents

#### Enzymes

PmeI (NEB R0560), PspOMI (NEB R0653), DNAse I (RNAse free) (NEB M0303)

#### Antibodies

GFAP (75-240, NeuroMab), MAP2 (AB5622, Millipore), ANTI-FLAG M2 antibody, Mouse monoclonal (mouse, Sigma F1804), alpha Tubulin Monoclonal Antibody (YL1/2) (rat, ThermoFisher MA1-80017), Anti-Puromycin Antibody, clone 12D10 (mouse, Sigma MABE343), GAPDH (Proteintech 60004-1-Ig), Actin (Sigma A1978), ATF-4 (SCBT sc-200), eIF1 (CST 12496S), eIF5 (D5G9) (CST 13894S), RPS11 (abcam ab157101), HRP goat anti-mouse (Jackson 115-035-146), HRP goat anti-rat (Jackson 112-035-003), Alexa Flour IR rat 800 (LiCOR 926-32219)

#### Kits/Other Reagents

HiScribe® T7 High Yield RNA Synthesis Kit (NEB E2040), 3′-O-Me- m^7^G(5’)ppp(5’)G RNA Cap Structure Analog (NEB S1411), m^7^G(5’)ppp(5’)A RNA Cap Structure Analog (NEB S1405), *E. coli* Poly(A) Polymerase kit (M0276), RNA Clean & Concentrator-25 kit (Zymo Research 1018), Lipofectamine MessengerMAX (Thermo Fisher, LMRNA008), Glolysis Buffer (Promega E26661), NanoGlo Substrate and Buffer (Promega N1150), One-Glo Solution (Promega E6130), CellTiter-Glo Solution (Promega G7570), Nano-Glo HiBiT Lytic Substrate and Buffer (Promega N3030), Complete Mini protease inhibitor tablet (Roche, 11836153001)

#### Drugs

Puromycin (Sigma P8833), Cycloheximide (Sigma 01810), Thapsigargin (Thermo T7459), Sodium Arsenite (Fluka 35000)

#### Cell Culture Reagents

DMEM with high glucose (Gibco 11965118), FBS (Bio-Techne S11150), 1% penicillin-streptomycin (Gibco 15070063), 1% MEM non-essential amino acids (Gibco 11140050), Neurobasal-A (Gibco 10888022), GlutaMax (Gibco 35050061), B27 Supplement (Gibco 17504044), mTeSR plus (Stem Cell, 100-0276), TeSR-E8 (Stemcell 05990), DMEM F12 (Gibco 11320033), Astrocyte Medium (ScienCell 1801)

## RESULTS

### Cell type-specific differences in RAN translation efficiency and regulation

To examine the mechanism of RAN translation initiation in neurons, we utilized a series of nanoluciferase (NL) reporters to assess translation efficiency across multiple systems (Fig. 1A). These included reporters for all three frames of translation from the expanded repeat in C9 ALS/FTD as well as a reporter for the poly-glycine protein produced across the expanded CGG repeat in the 5′ UTR of *FMR1* in FXTAS. As previously demonstrated (22, 25), nanoluciferase signal is abrogated by mutation of the nanoluciferase AUG start site codon to GGG but partially recovered by placement of repeats within their native sequence context 5′ to the repeat, implying that translation initiates either upstream of or within the repeats themselves. We generated *in vitro* transcribed mRNA from these reporters with a fully functional m^7^G cap and quantified translation following transfection into HEK293 cells, HeLa cells, or rat primary hippocampal neurons via the nanoluciferase assay. Consistent with previous results (22, 25), RAN translation was an inefficient process, with expression of C9 RAN reporters in all three reading frames, as well as the +1CGG100 FMR1 RAN reporter in FXTAS, at ∼0.5% to 3% of that observed from a canonical AUG-NL control in HEK293 cells (Fig S1A–D). For C9 RAN reporters, translation from the GA reading frame, in which the nanoluciferase tag is in frame with a CUG near cognate start codon, was 4-fold higher than that of GR and 8-fold higher than the GP reading frame in HEK293 cells (Fig. S1I). In HeLa cells and rat primary hippocampal neurons GP and GR production was also highly inefficient compared to GA; in neurons, GA production was 25-fold higher than GP and 27-fold higher than GR (Fig. S1J–K). This ratio is consistent with DPR abundance as measured in autopsied patient brains (38) and in other studies (53).

**Figure 1:**
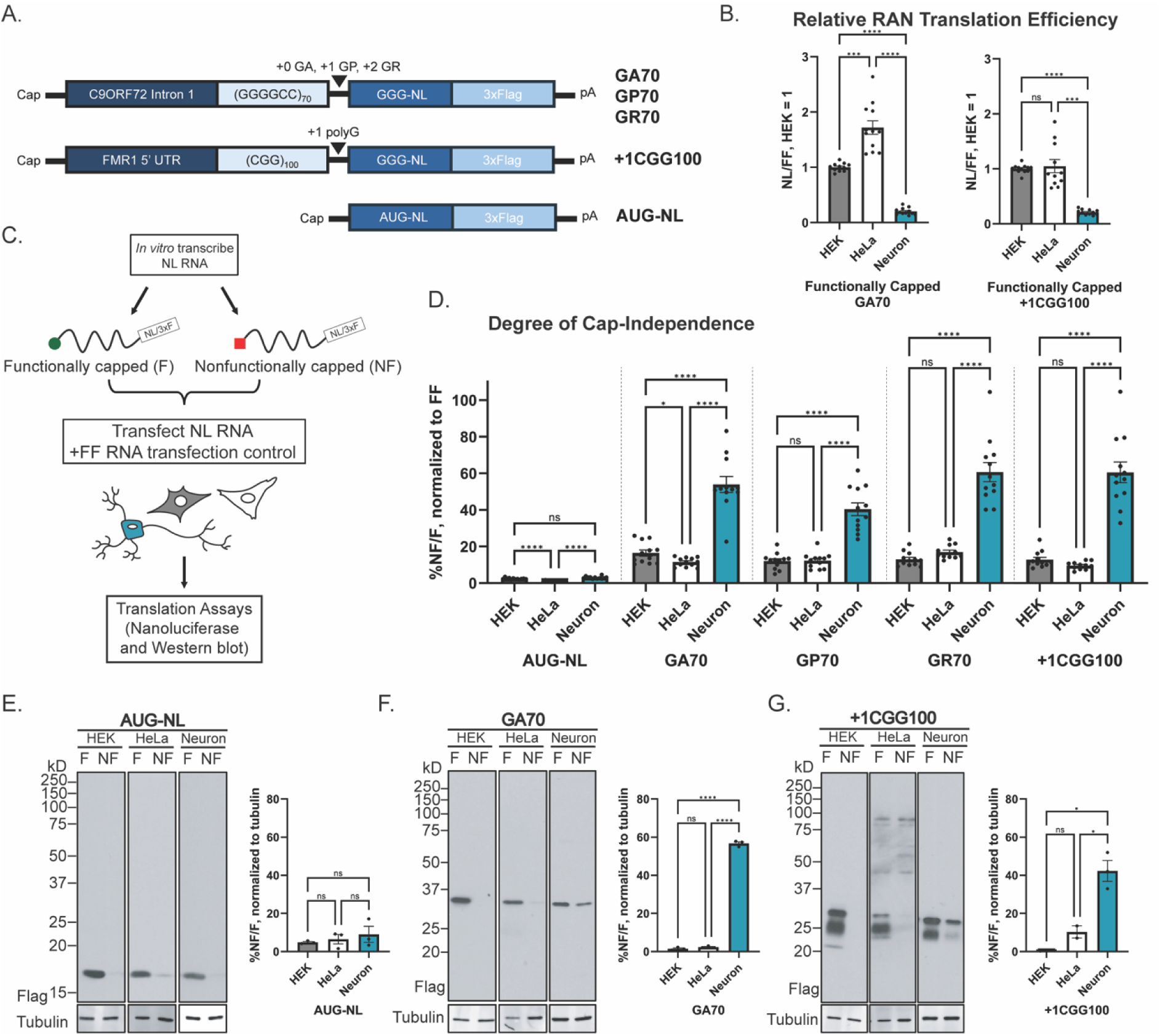
RAN translation in neurons is less efficient but more cap independent. (A) Schematic of linearized nanoluciferase (NL) reporter mRNAs. (B) Relative expression of RAN reporters in HEK293 cells, HeLa cells, and rat primary hippocampal neurons after transfection with functionally capped mRNA for 24 hours. Data is expressed as NL signal normalized to FF, n = 12 (C) Schematic of experimental strategy. (D) Degree of cap-independence of reporters after transfection into HEK293 cells, HeLa cells, and rat primary hippocampal neurons for 24 hours. Data is expressed as percent signal from nonfunctionally capped mRNA (NF) compared to the functionally capped mRNA (F), normalized to FF, n = 12. (E–G) Representative Western blots of HEK293 cells, HeLa cells, and rat primary hippocampal neurons transfected with functionally capped (F) and nonfunctionally capped (NF) mRNA reporters for 24 hours. Quantification data is expressed as percent intensity of the nonfunctionally capped mRNA (NF) compared to the functionally capped mRNA (F), normalized to tubulin (loading control), n = 3. Graphs represent mean ± SEM; Brown-Forsythe one-way ANOVA with Dunnet’s multiple comparison correction. *P < 0.05, **P<0.01, ***P < 0.001, ****P < 0.0001. GA = polyglycine-alanine, GP = polyglycine-proline, GR = polyglycine-arginine, +1CGG = polyglycine, FF = firefly.

The relative efficiency of RAN translation varied significantly across cellular models. Translation of both a GA reading frame C9 RAN reporter as well as the +1CGG100 FMR1 RAN reporter in FXTAS were 5-fold less efficient in rat primary hippocampal neurons compared to reporters for canonical translation than in HEK293 cells (Fig. 1B) (53). This same relationship was also observed for C9 RAN translation in the less efficient GP and GR reading frames (Fig. S1B–C).

To assess the translation efficiency of cap-independent RAN translation, we next *in vitro* transcribed mRNA from these reporters with either a functional cap (F) or a nonfunctional (NF) cap analogue that stabilizes the mRNA from degradation but does not bind to the initiation factor eIF4E to allow for mRNA activation (Fig. 1C). Expression of C9 RAN reporters in all three reading frames, as well as the +1CGG100 FMR1 RAN reporter, was well below expression from a canonical AUG-NL control with relatively restricted translation in neurons in both capping conditions (Fig. S1A–H). These data indicate that translation of these products with or without a functional cap remains highly inefficient compared to canonical translation.

To determine the degree of cap-independent translation in these cell types, we quantified the relative translation of functionally capped compared to nonfunctionally capped RNAs in different cell types.

As expected, translation of an AUG-NL control for canonical translation was highly cap-dependent in all cell types (Fig. 1D). Similarly, expression of all four different RAN reporters was largely cap-dependent in HEK293 and HeLa cells (Fig 1D). However, RAN translation exhibited a neuron-specific enhancement in the proportion of cap-independent translation (Fig. 1D) compared to cap-dependent translation. These results were confirmed via western blot of cells transfected with the same RNAs (Fig. 1E–G). As with capped mRNA reporters, RAN translation from nonfunctionally capped mRNA reporters was most efficient in the GA reading frame compared to the GP and GR reading frames across HEK293 cells, HeLa cells, and rat primary hippocampal neurons (Fig. S1 I–N). To assess if the high degree of cap-independence in neurons was due to their post-mitotic status, we treated HEK293 cells with hydroxurea, a cell-cycle inhibitor, and quantified the degree of cap-independence. A expected, Hydroxyurea-treated cells exhibited a lower degree of cap-independence than vehicle treated cells (Fig. S2), suggesting that the relative cap-independence in neurons cannot be fully explained by cell-cycle exit.

### Human-derived neurons exhibit efficient cap-independent RAN translation

To discern whether the observed difference in cap-independent translation in rat hippocampal neurons was a species-specific phenomenon, as well as to compare neuronal translation to a non-immortalized cell line, we differentiated healthy control iPSCs into cortical forebrain iNeurons or iAstrocytes and confirmed successful differentiation via immunocytochemistry for cell type specific markers microtubule associated protein 2 (MAP2) and glial fibrillary acidic protein (GFAP), respectively (Fig. 2A). After transfection of these different human cell types with functionally capped *in vitro* transcribed reporter mRNAs as described above, we found that RAN translation from both the GA C9 reporter and the +1CGG100 FMR1 RAN reporter were 44% and 32% as efficient, respectively, in iNeurons compared to iAstrocytes (Fig. 2B). RAN translation remained highly inefficient compared to expression of our canonical AUG-NL control, with efficiencies ranging from ∼0.01% to 2% AUG-NL signal (Fig. S3A–H). iNeurons showed low levels of relative cap-dependent RAN translation compared to iAstrocytes (Fig. S3A–D). However, nonfunctionally capped expression levels were similar between the cell types (Fig. S3E–H), indicating high relative levels of cap-independent RAN translation in the iNeurons compared to another differentiated neural cell type.

**Figure 2:**
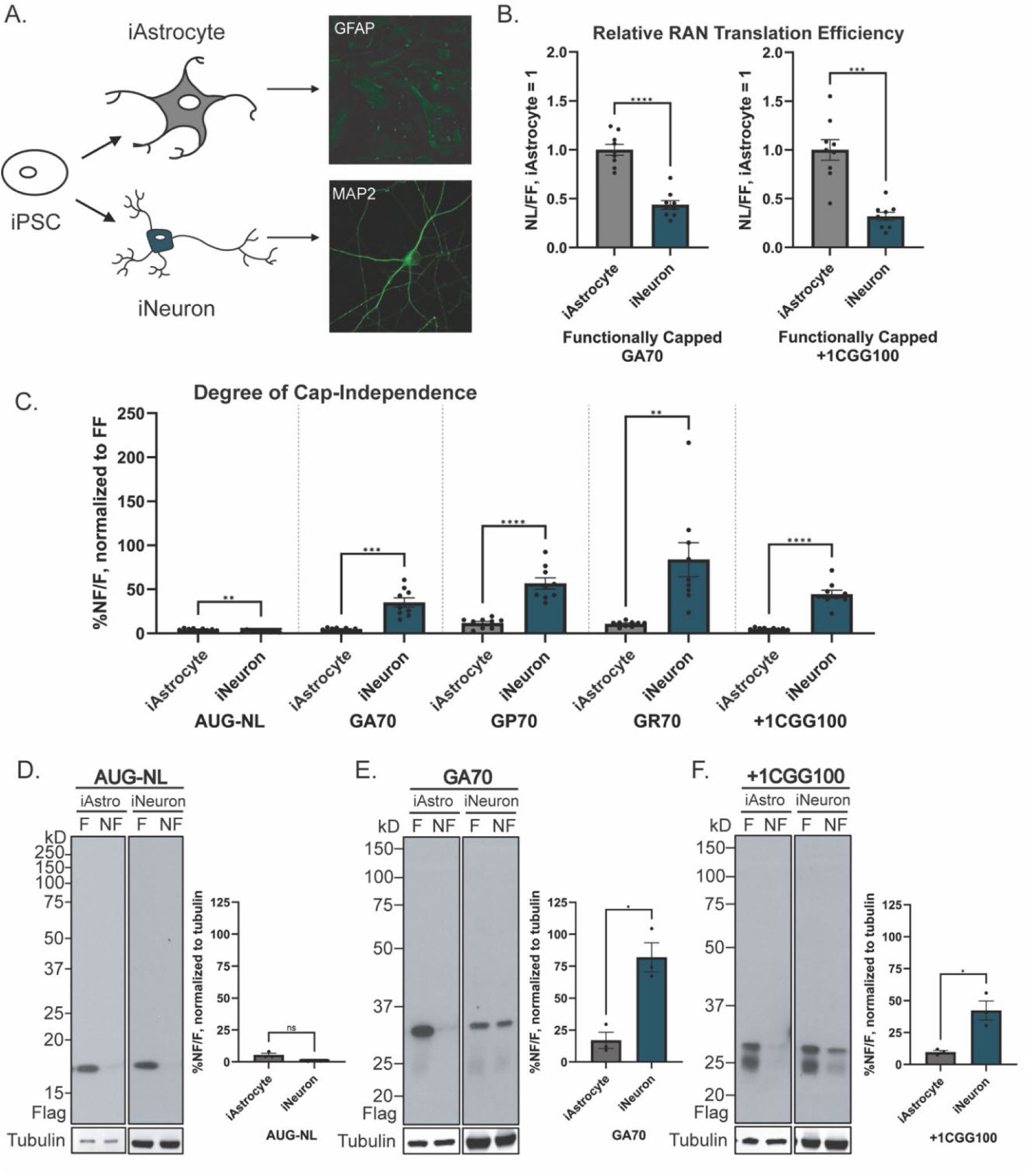
Cap-independent RAN translation is efficient in human-derived neurons. (A) Differentiation schema and representative ICC images for confirmation of differentiation of iAstrocytes and iNeurons. (B) Relative expression of RAN reporters in iAstrocytes and iNeurons after transfection with functionally capped mRNA for 24 hours. Data is expressed as NL signal normalized to FF, n = 9. (C) Degree of cap-independence of reporters after transfection into iAstrocytes and iNeurons for 24 hours. Data is expressed as percent signal in the nonfunctionally capped condition (NF) compared to the functionally capped condition (F), normalized to FF, n = 9. (D–F) Representative Western blots of iAstrocytes and iNeurons transfected with functionally capped (F) and nonfunctionally capped (NF) mRNA reporters for 24 hours and their quantification. Quantification data is expressed as percent intensity of the nonfunctionally capped mRNA (NF) compared to the functionally capped mRNA (F), normalized to tubulin (loading control), n = 3. Graphs represent mean ± SEM; Two-tailed Student’s t test with Welch’s correction. *P < 0.05, **P<0.01, ***P < 0.001, ****P < 0.0001.

To determine the degree of cap-independent translation in these cell types, we again quantified the translation efficiency ratio of nonfunctionally capped to functionally capped mRNA reporters. Translation of AUG-NL remained highly cap-dependent while all four RAN reporters demonstrated a high degree of cap-independent translation in the iNeurons but not in iAstrocytes derived from the same iPSC line (Fig. 2C). These results were confirmed via western blot of cells transfected with the same reporter mRNAs (Fig. 2D–F), indicating that a higher proportion of cap-independent translation is present in human neurons.

GA translation remained the most efficient across both iPSC-derived cell types, with GP and GR translation efficiencies ranging from 1 to 3% that of GA (Fig. S3I–L), indicating a continued preference for CUG-initiated translation for both functionally capped and nonfunctionally capped conditions.

Interestingly in iNeurons, we observed a greater degree of cap-independence in the GP and GR frame than in the GA frame (Fig. 2C). GP and GR are not in frame with the near-cognate CUG that drives efficient translation of GA. Instead, they are thought to be produced through either frameshifting or non-cognate initiation mechanisms (25, 26). Together, these data indicate that human iNeurons selectively favor cap-independent RAN translation in all three reading frames of C9 reporters and in the poly-glycine reading frame of FMR1 reporters.

### Inhibition of cap-binding initiation factor eIF4E preferentially enhances reporter-based and endogenous RAN translation in neurons

While utilizing in vitro transcribed mRNAs allows for precise modulation of cap-dependency in cells and the studies above using these reporters mimic relative translation levels seen in disease (38), the use of transiently transfected constructs may not fully recapitulate RAN translation from the endogenous locus in human neurons. To examine cap-independent initiation of endogenous C9-RAN translation, we utilized eFT-565, which competitively inhibits cap-binding factor eIF4E from binding to the 5’ cap of the mRNA to activate it and allow for recruitment of the 43S preinitiation complex to the ribosome (59, 60).

Treatment of HEK293 cells with eFT-565 inhibited global translation to a similar degree as cycloheximide, observed via measurement of puromycin incorporation with a surface sensing of translation (SUnSET) assay (Fig. 3A–B) (58). eFT-565 treatment induced minimal toxicity in HEK293 cells and rat primary hippocampal neurons (Fig. S4A–B), while inhibiting translation efficiency of both a canonical translation control AUG-NL reporter and a firefly luciferase reporter in both cell types (Fig. 3C–D, Fig. S4C–D). In HEK293 cells transfected with a functionally capped GA70 reporter, eFT-565 treatment decreased GA production (Fig. 3E), further supporting its cap-dependence in this cell type. However, eFT-565 treatment of rat primary hippocampal neurons transfected with the same mRNA reporter paradoxically increased GA frame RAN translation (Fig. 3F), providing orthogonal evidence of cell-type specific differences in RAN translational cap-dependency. Interestingly, eFT-565 treatment increased expression of a nonfunctionally capped GA70 reporter in both HEK293 cells and in rat neurons (Fig. S4E–F). For GP frame production we observed similar results as in the GA frame: eFT-565 treatment strongly inhibited expression of a functionally capped GP70 reporter in HEK293 cells, with differential effects in neurons (Fig. S4G–H).

**Figure 3:**
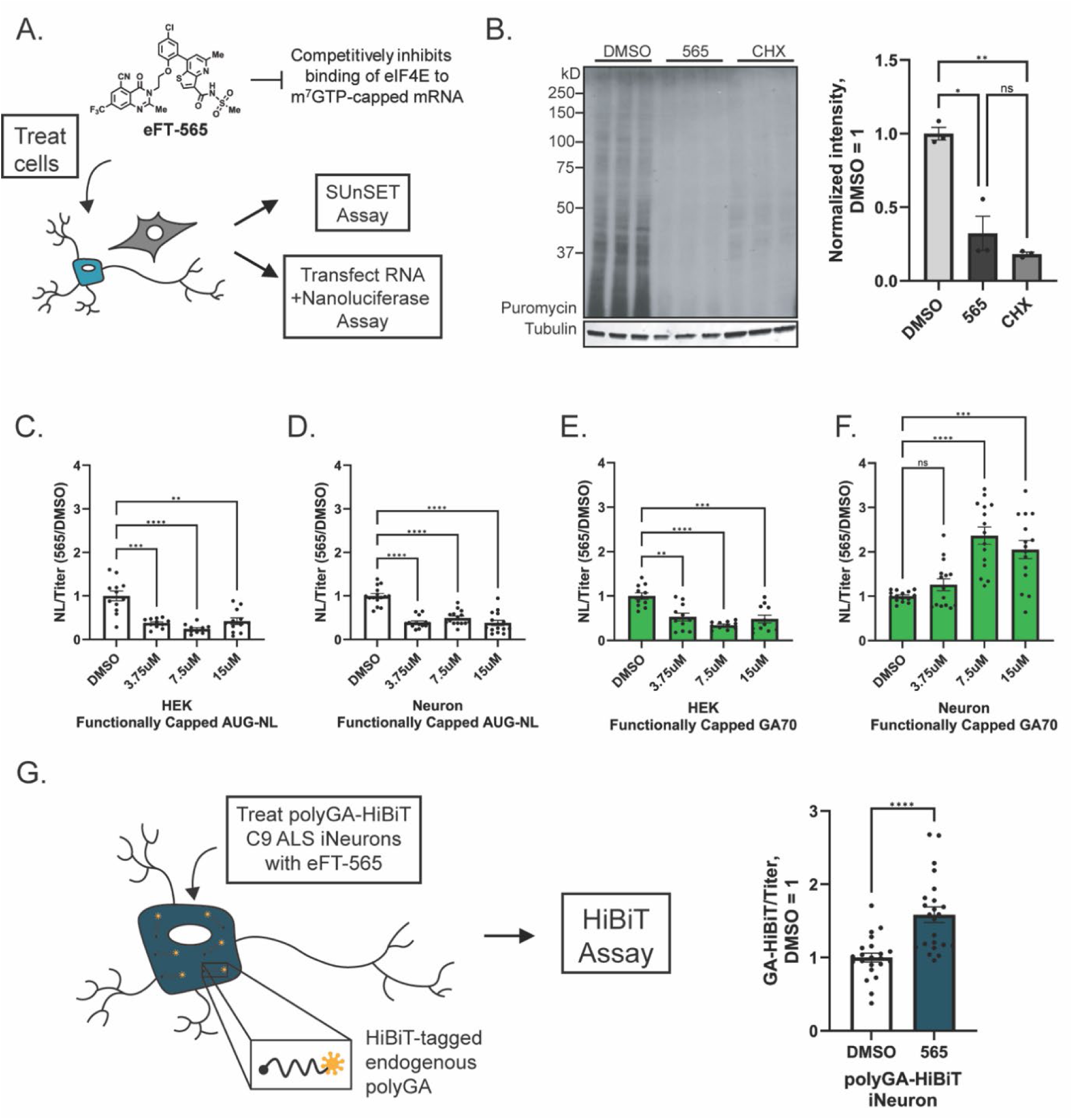
Inhibiting cap-binding factor eIF4E selectively enhances neuronal RAN translation. (A) Schematic of experimental strategy. (B) SUnSET assay of puromycin incorporation in HEK293 cells treated with 24 hours of DMSO, 24 hours of 7.5μM eIF4E competitive inhibitor eFT-565 (565), or 15 minutes of 100μg/mL cycloheximide (CHX) as a positive control. Data is expressed as intensity in each condition normalized to tubulin (loading control) and relative to DMSO treated condition, n = 3. (C–F) Relative response to treatment with eFT-565. HEK293 cells (C/E) and rat primary hippocampal neurons (D/F) were treated with 0 – 15μM eFT-565, transfected with functionally capped AUG (C–D) or GA70 (E–F) reporters 4.5 hours later, and assayed for translation 19.5 hours later. Data is expressed as NL signal normalized to cell titer and relative to the DMSO treated condition, n = 12-15. (G) Endogenous PolyGA-HibiT levels in polyGA-HiBiT tagged iNeurons after 24 hours of treatment with DMSO or 7.5μM eFT-565. Data expressed as HiBiT signal normalized to cell titer and relative to DMSO treated condition, n = 23. Graphs represent mean ± SEM; (B–F) Brown-Forsythe one-way ANOVA with Dunnet’s multiple comparison correction. (G) Two-tailed Student’s t test with Welch’s correction. *P < 0.05, **P<0.01, ***P < 0.001, ****P < 0.0001.

To assess the impact of eIF4E inhibition on endogenous C9-RAN translation, we utilized a recently generated patient-derived iPSC cell line in which CRISPR-Cas9 was used to insert a HiBiT tag downstream of the polyGA frame in the C9orf72 gene of a patient with C9ALS/FTD for the quantification of endogenous GA (61). This line harbors approximately 700 GGGGCC repeats (61), allowing for examination of much larger repeats than are technically feasible with reporters. eFT-565 treatment of GA-HiBiT iNeurons robustly enhanced endogenous GA frame translation (Fig. 3G), supporting results observed in exogenously introduced reporter-based assays. This finding indicates that endogenous translation of the GA frame RAN product in human neurons can be initiated in a cap-independent manner and its total translation is enhanced under cap-inhibited conditions.

### Cap-independent RAN translation resists inhibition by the integrated stress response

The integrated stress response (ISR) maintains cellular homeostasis through the phosphorylation of eIF2α at serine 51 (Fig. 4A) (62, 63). This phosphorylation inhibits global cap-dependent translation and allows translation of specific mRNAs as an adaptive response (Fig. 4A) (62, 63). Dysregulation and overload of this response are observed in many neurodegenerative disorders, including ALS/FTD, leading to cell death (63). Activation of the ISR upregulates RAN translation, leading to a feed-forward loop of disease pathogenesis in which DPRs trigger the ISR and activation of the ISR further upregulates production of the neurotoxic DPRs (25, 31, 32, 53). We therefore assessed whether cap- independent RAN translation is responsive to ISR activation in different cell types.

**Figure 4:**
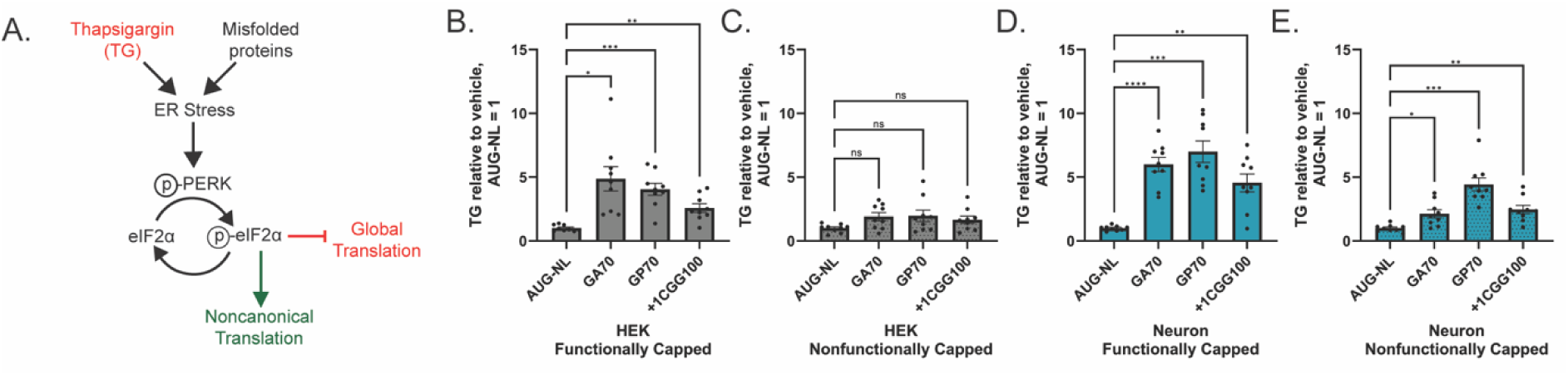
Neuronal cap-independent RAN translation is resistant to integrated stress response activation. (A) Schematic of the ER stress branch of the integrated stress response. (B–E) Relative expression of reporters in HEK293 cells (B–C) and rat primary hippocampal neurons (D–E) treated with 2μM thapsigargin, transfected with functionally capped (B/D) or nonfunctionally capped (C/E) reporters, and assayed for translation 5 hours later. Data is expressed as ratio of NL signal normalized to titer and relative to AUG response, n = 9. Graphs represent mean ± SEM; Brown-Forsythe one-way ANOVA with Dunnet’s multiple comparison correction. *P<0.05, **P<0.01, ***P<0.001, ****P<0.0001.

HEK293 cells and rat primary hippocampal neurons treated with the ER calcium pump inhibitor thapsigargin (TG) to trigger ER stress (Fig. 4A) were transduced with functionally capped or nonfunctionally capped RAN translation and control reporter mRNAs (see methods). RAN translation of both functionally capped C9 and FMR1 reporters was resistant to TG treatment compared to canonical AUG-NL control reporters in both HEK293 and hippocampal neurons (Fig. 4B/D). Cap-independent RAN translation was modestly resistant to TG treatment in neurons (Fig. 4E) but was not significantly resistant in HEK293 cells (Fig. 4C). Treatment with sodium arsenite, which triggers the oxidative stress branch of the ISR through the kinase HRI (Fig. S5A), produced similar results in rat primary hippocampal neurons, where both functionally capped and non-functionally capped C9 RAN reporter RNAs were resistant to arsenite treatment as compared to a canonical AUG-NL control (Fig. S5D–E). Both cell types exhibited similar ISR activation in response to these stressors (Fig S45–G).

Together, these data indicate that both cap-dependent and cap-independent RAN translation are resistant to ISR activation in neurons, although cap-dependent RAN translation exhibited a greater degree of resistance.

### CUG near cognate codon mutation favors cap-independent RAN translation

GA frame C9-RAN translation is largely dependent on a near cognate CUG codon located 24 nucleotides upstream of the repeat in a strong Kozak sequence for initiation (14, 25, 26, 28, 32). It is unclear if cap-independent C9-RAN translation utilizes this codon as cap-independent, IRES mediated-translation in other contexts can occur in the presence or absence of ribosomal scanning and with less stringent requirements for start codon identity (36).

The relatively high efficiency of GA production from both functionally capped and nonfunctionally capped C9 reporters across cell types (Figs. S1I–N, S3I–L) suggests that cap-independent GA frame C9-RAN translation likely also utilizes this CUG codon. To examine this question directly, we utilized a GA70 reporter with the CUG codon mutated to a CCC non-cognate codon (Fig. 5A). This CCC- mutation inhibited GA RAN translation in all contexts (Fig. 5B–C), indicating that both cap-dependent and cap-independent GA frame translation utilize this codon. However, the cap-independence of the CCC-mutated GA70 reporter was significantly greater than that of its non-mutated CUG GA70 counterpart (Fig. 5D), suggesting that the CCC-mutated GA70 reporter favors cap-independent RAN translation. This observation was most robust in rat primary hippocampal neurons, where the degree of cap-independence nearly doubled with the mutation (Fig. 5D).

**Figure 5:**
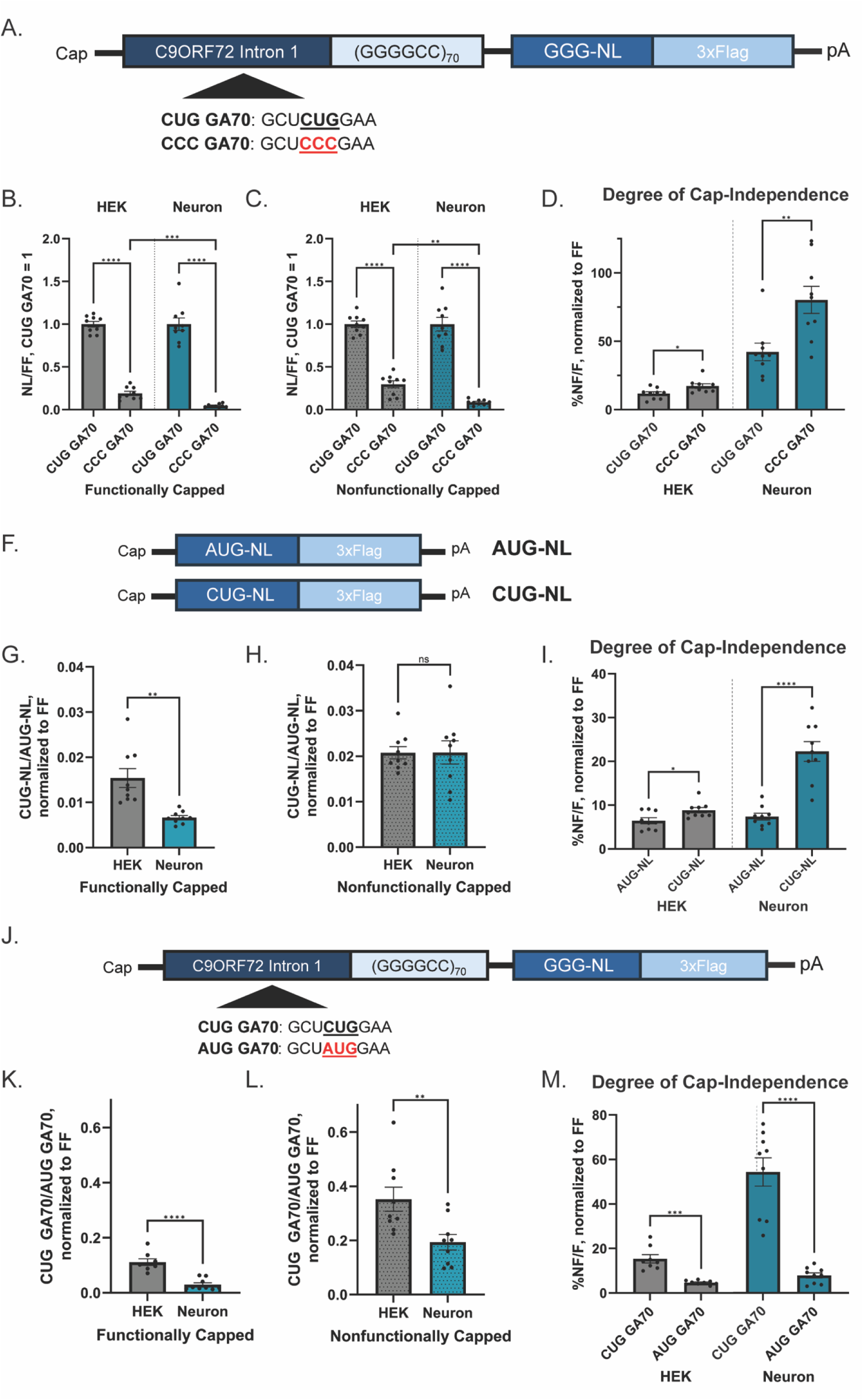
Altered start codon stringency in neurons drives greater efficiency of cap-independent RAN translation. (A) Schematic of linearized NL reporter mRNAs. (B–C) Relative expression of CCC GA70 in HEK293 cells and rat primary hippocampal neurons transfected with functionally capped (B) or nonfunctionally capped (C) mRNA for 24 hours. Data is expressed as NL signal normalized to FF and relative to CUG GA70, n = 9. (D) Degree of cap-independence of reporters after transfection into HEK293 cells and rat primary hippocampal neurons for 24 hours. Data is expressed as percent NL signal in the nonfunctionally capped condition (NF) compared to the functionally capped condition (F), normalized to FF, n = 9. (F) Schematic of linearized non-repeat NL reporter mRNAs. (G–H) Ratio of CUG-NL/AUG-NL in HEK293 cells and rat primary hippocampal neurons transfected with functionally capped (G) or nonfunctionally capped (H) mRNA for 24 hours. Data is expressed as CUG-NL signal compared to AUG-NL signal normalized to FF, n = 9. (I) Degree of cap-independence of reporters after transfection into HEK293 cells and rat primary hippocampal neurons for 24 hours. Data is expressed as percent NL signal in the nonfunctionally capped condition (NF) compared to the functionally capped condition (F), normalized to FF, n = 9. (J) Schematic of linearized NL reporter mRNAs. (K–L) Ratio of expression of CUG GA70/AUG GA70 in HEK293 cells and rat primary hippocampal neurons transfected with functionally capped (F) or nonfunctionally capped (NF) mRNA for 24 hours. Data is expressed as CUG GA70 signal compared to AUG GA70 signal normalized to FF, n =9. (M) Degree of cap-independence of reporters after transfection into HEK293 cells and rat primary hippocampal neurons for 24 hours. Data is expressed as percent NL signal in the nonfunctionally capped condition (NF) compared to the functionally capped condition (F), normalized to FF, n = 9. Graphs represent mean ± SEM; (B–C) Brown-Forsythe one-way ANOVA with Dunnet’s multiple comparison correction; (D, G–I, K–L) Two-tailed Student’s t test with Welch’s correction. *P < 0.05, **P<0.01, ***P < 0.001, ****P < 0.0001.

### Neurons exhibit high degree of start codon stringency

Mutation of this CUG near-cognate codon affected RAN translation efficiency in rat primary hippocampal neurons to a much greater degree compared to HEK293 cells (Fig 5B–C). This finding suggested that neurons might exhibit a greater degree of start codon fidelity compared to other cell types. Start codon recognition is mediated by multiple factors, including a complex and adaptive interplay between eIF1 and eIF5 (9, 15). eIF1 enhances start codon stringency through mediation of AUG recognition, while eIF5 enhances initiation at non-AUG start sites (9, 15). Dynamic changes to their ratio of expression can modulate start site fidelity across different contexts and modulates RAN translation (14, 15, 64).

To directly examine the degree of start codon stringency across cell types, we utilized nanoluciferase reporters initiated by either an AUG start site (AUG-NL) or by a CUG (CUG-NL) without any repeat or C9orf72 relevant sequence context (Fig. 5F). Transfecting mRNA of these reporters into rat primary hippocampal neurons or HEK293 cells revealed low levels of CUG-initiated translation in both cell types (Fig. 5G–H). CUG-initiated translation from the functionally capped reporters in neurons was 42% as efficient as in the HEK293 cells (Fig. 5G), indicating a higher degree of start codon stringency in the neurons for cap-dependent translation. This high degree of start codon-stringency was seen with other near-cognate reporters as well in both human and rat neurons (Fig. S6). Interestingly, this cell-type specific effect was not seen with nonfunctionally capped RNA: there was no difference observed in the CUG/AUG ratio between the two cell types (Fig. 5H). Overall, AUG-NL translation exhibited a greater degree of cap-dependence than CUG-NL, especially in neurons (Fig. 5I). Together, these results indicate that start codon stringency for nanoluciferase reporters is higher in neurons compared to HEK293 cells.

### Neuronal start codon stringency favors cap-independent RAN translation

To examine if CUG-initiated C9-RAN translation is similarly restricted in neurons, we utilized GA70 reporters with either the native CUG codon or with an AUG codon substituted into its place (Fig. 5J). As previously shown (25), AUG-initiated translation through the repeat was more efficient than CUG-initiated RAN translation (Fig. 5K–L) and this enhanced efficiency was observed for both functionally capped and non-functionally capped mRNAs and in both neurons and in HEK293 cells (Fig. 5K–L).

However, AUG-GA70 reporters exhibited a higher cap-dependence compared to the native CUG- GA70 RAN translation reporters. This effect was particularly pronounced in neurons (Fig. 5M).

### Initiation factor eIF1 exhibits cell-type specific localization patterns

Global translation levels in both human iNeurons (Fig. 6A) and rat primary hippocampal neurons (Fig. S7A) were lower than their comparator cells via SUnSET assay. To interrogate the mechanism by which neuronal start codon stringency is enhanced, we first examined initiation factor expression. Via transcriptomic analysis, we found that neurons express higher levels of eIF1, 5MP, and eIF1A transcripts while there was no significant difference in eIF5, eIF5B, or RPS111 ribosomal protein transcript levels (Fig. S7B). In contrast, compared to iAstrocytes, iNeurons exhibit lower levels of both ribosomal marker RPS11 and, surprisingly, initiation factor eIF1, which enhances start codon stringency (Fig. 6B). eIF5 expression was not significantly different between the cell types (Fig. 6B).

**Figure 6:**
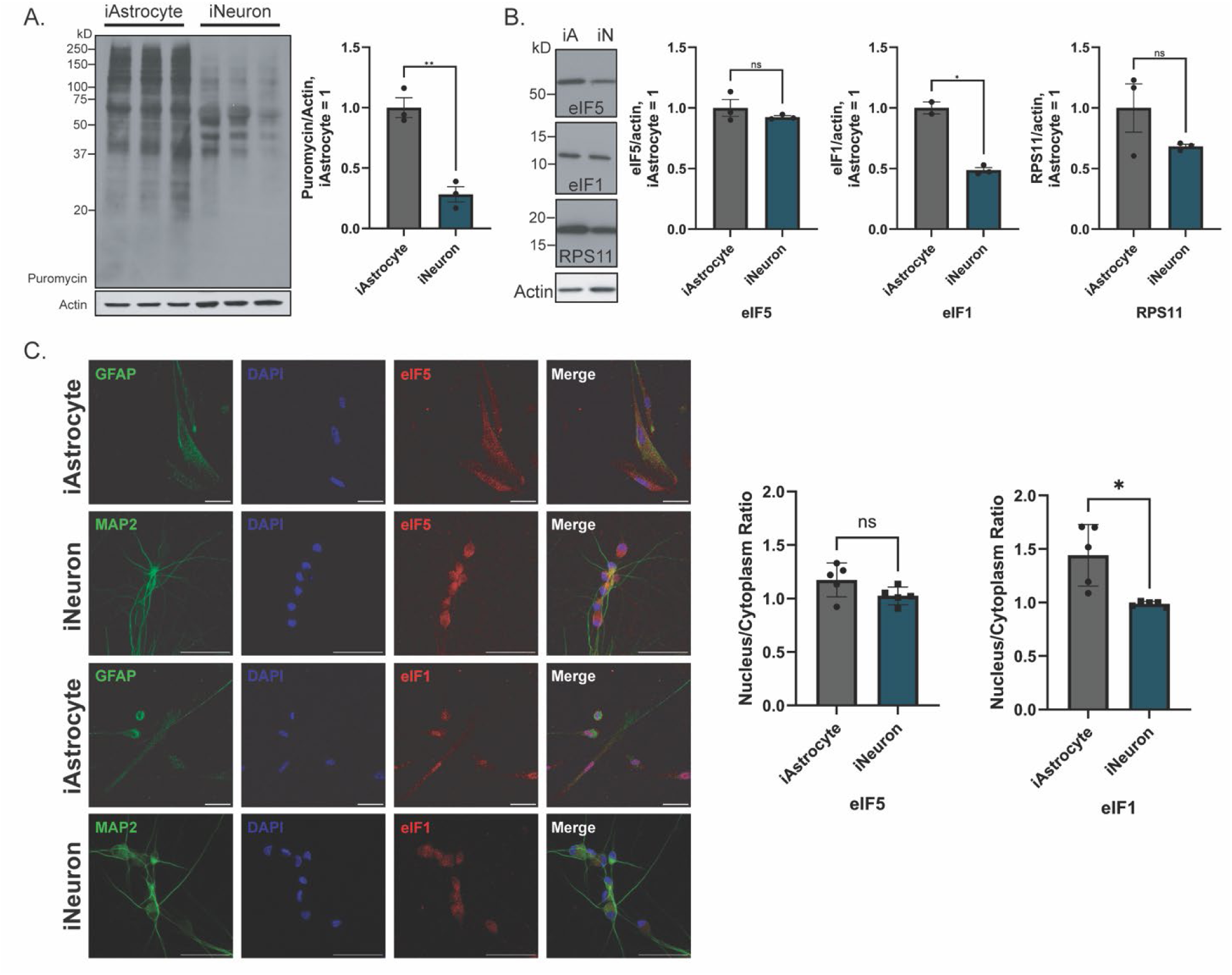
Global Translation rates and initiation factor eIF1 localization are cell-type specific. (A) SUnSET assay and quantification of puromycin incorporation in iAstrocytes and iNeurons. Data is expressed as intensity in each condition normalized to actin (loading control), n = 3. (B) Representative Western blots and quantification of iAstrocytes and iNeurons probed for initiation factors and ribosomal marker. Data is expressed as intensity in each condition normalized to actin, n = 3. (C) Representative ICC images and quantification of nuclear to cytoplasmic ratio of eIF1 and eIF5 in iAstrocytes and iNeurons. Scale bar = 50µm. Data is expressed as ratio of nuclear and cytoplasmic brightness of all cells in a single image with each n representing an separate image, n = 5. Graphs represent mean ± SEM; two-tailed Student’s t test with Welch’s correction. *P < 0.05, **P<0.01, ***P < 0.001, ****P < 0.0001.

Next, we examined the nuclear to cytoplasmic ratio of eIF1 and eIF5 via ICC (Fig. 6C). We found a significantly higher cytoplasmic distribution of eIF1 in iNeurons than in iAstrocytes. In contrast, eIF5 distribution was not significantly different between the cell types (Fig. 6C).

### Start codon fidelity alters RAN translation

These data suggest a relationship between altered start codon fidelity, global translation rates and the reduced cap dependency observed for RAN translation in neurons. To assess this question more directly, we asked if we could reverse the observed lowering of cap-dependency for RAN translation in neurons by artificially loosening start codon stringency through overexpression of eIF5 (Fig. 7A). eIF5 is known to decrease start codon stringency (15) and increase both GA frame C9-RAN translation and CGG RAN translation (14, 64). Consistent with this prior work, overexpression of eIF5 specifically increased GA frame RAN translation from functionally capped and nonfunctionally capped mRNA in HEK293 cells (Fig. 7B–C). However, eIF5 overexpression had no effect on the degree of cap-dependence in these cells, which remained high (Fig. 7D). In neurons, overexpression enhanced translation from the functionally capped RAN translational reporter by almost 5 times the vehicle (Fig. 7E) but did not significantly enhance cap-independent RAN translation (Fig. 7F), indicating a specific effect on cap-dependent RAN translation. This effect of eIF5 overexpression was restricted to RAN translation and did not impact the cap-dependence of a canonically translated AUG-NL reporter (Fig. 7G). Together, these data support a direct relationship between higher start codon stringency and lower cap-dependent RAN translation in neurons.

**Figure 7:**
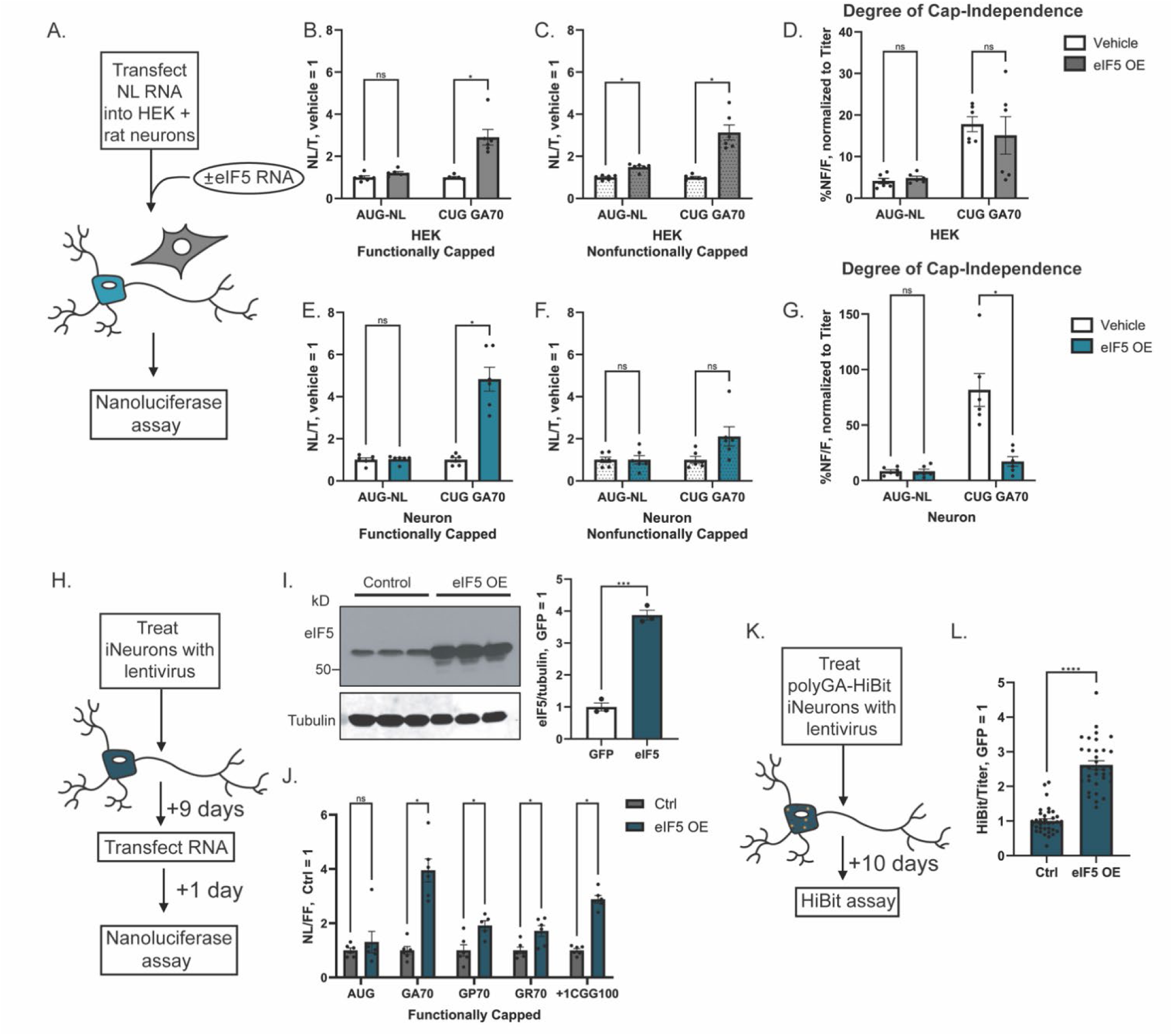
eIF5 overexpression alters degree of cap independence and RAN translation. (A) Schematic of experimental strategy for eIF5 overexpression via RNA co-transfection. (B–C) Relative expression of functionally capped (B) or nonfunctionally capped (C) reporters co-transfected with vehicle control or eIF5 in HEK293 cells for 24 hours. Data is expressed as NL signal normalized to titer and relative to vehicle control, n = 6. (D) Degree of cap-independence of reporters after transfection. Data is expressed as percent NL signal in the nonfunctionally capped condition (NF) compared to the functionally capped condition (F), normalized to titer, n = 6. (E–F) Relative expression of functionally capped (E) or nonfunctionally capped (F) reporters co-transfected with vehicle control or eif5 in rat primary hippocampal neurons for 24 hours. Data is expressed as NL signal normalized to titer and relative to vehicle control, n = 6. (G) Degree of cap-independence of reporters after transfection. Data is expressed as percent NL signal in the nonfunctionally capped condition compared to the functionally capped condition, normalized to titer, n = 6. (H) Schematic of experimental strategy for eIF5 overexpression via lentivirus. (I) Western blot and quantification of eIF5 levels after treatment with control (GFP) or eIF5 overexpression lentivirus. (J) Relative expression of functionally capped RNA reporters into iNeurons treated with control or eIF5 overexpression lentivirus. Data is expressed as NL signal normalized to FF with control set to 1, n = 6. (K) Schematic of experimental strategy for eIF5 overexpression via lentivirus into poly-GA HiBit iNeurons. (L) Endogenous PolyGA-HibiT levels in polyGA-HiBiT tagged iNeurons after 10 days of treatment with control or eIF5 overexpression lentivirus. Data expressed as HiBiT signal normalized to cell titer with control set to 1, n = 36. Graphs represent mean ± SEM; (B–G) Brown-Forsythe one-way ANOVA with Dunnet’s multiple comparison correction; (I, L) Two-tailed Student’s t test with Welch’s correction.; (J) Multiple Holm–Šídák’s two-tailed Student’s t test with Welch’s correction *P < 0.05, **P<0.01, ***P < 0.001, ****P < 0.0001.

To examine the effects of loosening start codon stringency on general RAN translation, we treated iNeurons with a lentivirus that overexpressed eIF5 (Fig. 7H) and confirmed its expression (Fig. 7I). In iNeurons, eIF5 overexpression enhanced translation from functionally capped C9 and +1CGG100 FMR1 RAN reporters after RNA transfection (Fig. 7J). To assess the effects of eIF5 overexpression on endogenous RAN translation, we overexpressed eIF5 in polyGA-HiBit tagged iNeurons using a lentivirus (Fig. 7K). This resulted in a 2.5 times increase in endogenous GA production (Fig. 7L).

## DISCUSSION

The pathogenic mechanisms that drive neuro-specific degeneration in nucleotide repeat expansion disorders remain enigmatic. RAN translation generates neurotoxic peptides that contribute to neurodegeneration. Prior studies found that RAN translation is highly cap-dependent in transfected cells, making the contribution of cap-independent RAN translation to disease pathogenesis uncertain (25, 26, 28, 31, 32, 53). This study examined RAN translation in both C9 ALS/FTD and FXTAS using both reporter-based assays and endogenous readouts to delineate cell-type specific differences in its regulation. We show that RAN translation in neurons is less efficient overall but also less dependent on the 5′ m^7^G cap. The relative inefficiency of RAN translation in neurons correlates with higher levels of start codon stringency that largely spares cap-independent RAN translation, driving a shift towards this initiation mechanism. These findings provide insight into neuronal translational regulation and inform our understanding of the mechanisms by which RAN translation might contribute to neurodegeneration in nucleotide repeat expansion disorders.

Supporting findings from Westergard et al. (53), we find that RAN translation is less efficient in neurons compared to other cell types (Figs. 1B, 2B, S1, S2). We uncovered a high degree of start codon stringency in neurons as a potential explanation for this relative inefficiency (Fig 5, S6). As an immortalized cell line, it is perhaps unsurprising that HEK293 cells demonstrate a low degree of start codon stringency given that cancer cells exhibit dysregulated translation control that favors non-AUG initiation events (65–68). However, given that cap-independence was also low in iAstrocytes (Figs. 1D, 2C), this finding is likely not to be specific to immortalized cells.

Start codon stringency is governed in part by the dynamic interaction of human eukaryotic initiation factors eIF1 and eIF5 (15). These two factors are part of a complicated auto- and cross-regulatory feedback loop (69) and compete for an overlapping binding site (15, 70). eIF1 enhances start codon stringency, encouraging proper recognition of the start codon in the ribosomal P site, and decreases general RAN translation, while eIF5 activity increases non-AUG initiation, including in RAN translation (9, 14, 15, 35, 64, 71). The abundance of these factors (72) did not explain the higher level of start codon stringency seen in neurons (Figs 6B, S7B). Overall protein expression of eIF1 in iNeurons was actually lower than in iAstrocytes (Fig 6B), indicating that overall levels of this initiation factor are not a clear driver of increased start codon stringency. However, we observed that eIF1 is selectively distributed into the cytoplasm in neurons (Fig 6). This finding builds off of previous work demonstrating that release of eIF1 from the nucleus has strong inhibitory effects during cell division on general translation but allows select RNAs to escape such suppression (73). Consistent with this finding, overexpression of eIF5 markedly enhances RAN translation and reverses the cap-independence of RAN translation observed in neurons (Fig 7). Thus, our data suggests that cytoplasmic redistribution of eIF1 could play roles in setting start codon stringency in contexts outside of cell division, with potentially broad implications for translational control in neurons and other cell types. However, our data does not preclude additional roles for other initiation factors such as eIF5MP (74) in neuronal start codon stringency or cap-independent RAN translation. Future studies will be needed to parse the relative contributions of such factors to this enigmatic process.

How cap-independent RAN translation works mechanistically is not well understood. Cap-independent translation in other contexts often occurs through the utilization of a typically highly structured internal ribosome entry site (IRES) that serves as a mechanism to recruit the ribosome to an mRNA while bypassing some key regulatory elements that control canonical translational initiation (36, 37, 75, 76). GC-rich repeat regions in C9 ALS/FTD and FXTAS form secondary structures such as hairpins and G-quadruplexes, which could potentially serve as IRESes and bypass the need for cap-binding. Indeed, GGGGCC repeats can bind to the 40S small ribosomal subunit in isolation (26, 77–79).

Our study indicates that cap-independent C9-RAN translation in the GA reading frame still prefers the use of a CUG initiation codon (Fig. 5A–D). This finding suggests that the repeat may act like a type I or II IRES site, offering ribosomal stabilization or recruitment in the absence of cap-binding to allow for scanning and use of this CUG codon. However, nonfunctionally capped CCC-mutant GA70 reporters are still translated at an efficiency of around 80% of their functionally capped counterparts in neurons, indicating that cap-independent translation may be initiated independent of the near-cognate codon in neurons - likely within the repeat itself (Fig. 5A–D). Additionally, CUG-initiated translation still occurs, albeit inefficiently, in the absence of any repeat or mRNA cap (Fig. 5H) as previously shown (69, 80). These findings are reminiscent of prior data on CGG repeats in the FMR1 locus where mutation of all upstream near-cognate codons reduced but did not eliminate FMRpolyG production (35). In HEK293T cells, RAN translation from a mutated FMR1 5′UTR remained dependent on eIF4A function (as it was inhibited by hippuristanol), was suppressed by overexpression of eIF1, and remained largely cap-dependent (35). Further studies will be needed to define the exact mechanism(s) of cap-independent RAN translation, but the data here suggests that it is likely to occur at differing levels of efficiency that are dependent on cell type and state.

While this study connects the high degree of cap-independent GA frame C9-RAN translation with a greater degree of start codon stringency in neurons, high neuronal cap-independence is also observed in the GP and GR reading frames (Figs. 1D, 2C) and their translation is also less efficient overall in neurons (Figs. S1, S2). GP and GR are out of frame with the CUG codon but are thought to in part be produced by frameshifting following GA frame initiation (25, 26), which would still be dependent on the CUG near cognate codon. However, these constructs exhibit varying levels of dependence on the presence of this CUG codon dependent on cellular context (25, 26). As such, start codon stringency likely does not fully explain neuronal regulation of their production.

While start codon stringency provides a potential explanation for the neuronal behavior of RAN translation, this manuscript has several limitations. It remains possible that variation in the expression levels of other specific RAN translational modulators or ITAFs in neurons could explain some parts of our observations. These factors could be especially important for GP and GR frame C9-RAN translation which are less dependent on a near-cognate codon for initiation. Additionally, this study relies heavily on reporter-based assays. Although the efficiency of these reporters mirrors the relative accumulation of DPR products in patient brains (Figs. S1I–N, S2I–L), due to technical restrictions the expansions in these reporters are limited to 70 repeats, far below the hundreds to thousands typically seen in patients (1, 4, 7, 81). Secondary structure is particularly important for IRES-mediated translation, so this difference could be especially salient when studying cap-independent RAN translation. This makes our studies using a HiBiT tag to monitor the effect of eIF4E inhibition on endogenous GA RAN translation from a locus with approximately 700 repeats in patient-derived neurons (61) particularly important (Fig. 3G), as they indicate that endogenous GA production in neurons is enhanced under conditions that favor cap-independent translation.

If RAN translation in neurons drives disease pathogenesis in repeat expansion disorders, then understanding neuron-specific regulation of this process is an important step forward in therapy development for these currently untreatable diseases. While these processes may be inefficient, work here shows they are resilient to mechanisms which typically limit translation, making targeting them possibly important for therapeutic development. This study provides insights into both neuron specific mechanisms of RAN translation and neuronal translational regulation, uncovering a high degree of start codon stringency and cell-type specific modifications of the cap-dependence of RAN translation. These findings thus have implications for understanding the synthesis of toxic RAN translation products across multiple diseases and establish a baseline upon which we can better understand RAN translational modifiers as potential therapeutic targets.

## ACKNOWLEDGEMENTS

We thank Jesse Duque and Jason C. Rech, PhD for their work in the development, testing, and production of eFT-565. We thank Rachel O’Rourke for her assistance in obtaining eFT-565.

## AUTHOR CONTRIBUTIONS

Clare M. Wieland (Conceptualization [equal], Formal analysis [lead], Investigation [lead], Methodology [lead], Project administration [equal], Resources [lead], Visualization [lead], Writing – original draft [lead]); Shannon E. Wright (Conceptualization [supporting], Investigation [supporting], Methodology [supporting]); Sydney Willey (Investigation [supporting], Writing – original draft [supporting], Visualization [supporting]); Ishita Purwar (Formal analysis [supporting], Investigation [supporting], Writing – original draft [supporting]); Samantha J. Grudzien (Formal analysis [supporting], Investigation [supporting], Writing – original draft [supporting]); Amy Krans (Investigation [supporting], Methodology [supporting], Resources [supporting]); Erinn Laimon (Methodology [supporting], Resources [supporting], Writing – original draft [supporting]); Melissa J. Asher (Methodology [supporting], Resources [supporting]); Adrian Isaacs (Resources [supporting]); Amanda L. Garner (Resources [supporting]); Peter K. Todd (Conceptualization [equal], Funding acquisition [lead], Project administration [equal], Supervision [lead], Writing-review & editing [lead]).

## SUPPLEMENTARY DATA

Supplementary Data are available at NAR online.

## DATA AVAILABILITY

All study data are available in the article and/or the supplementary information.

## SUPPLEMENTAL FIGURES

**Figure S1:**
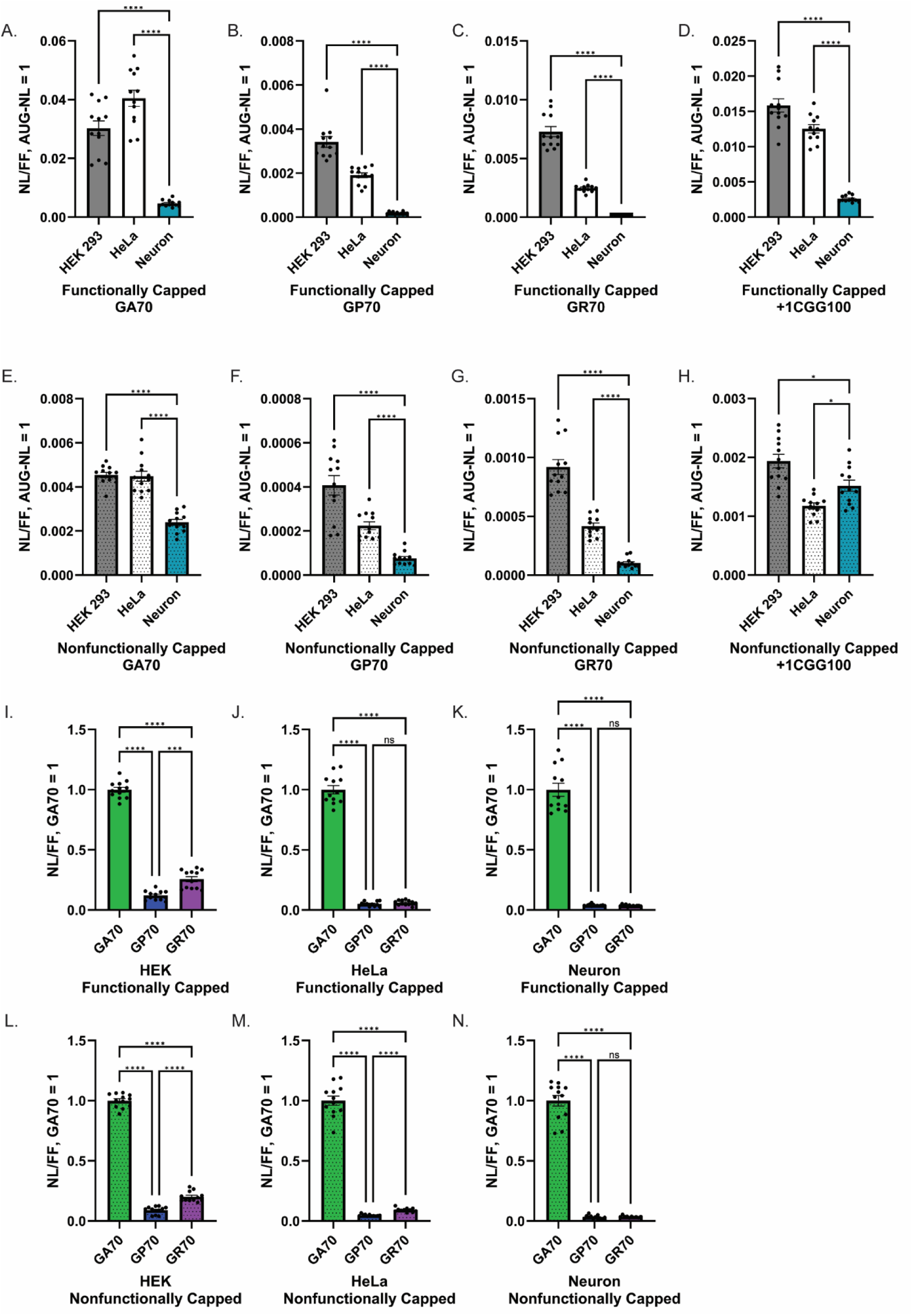
Expression of RAN reporters differ by cell type. (A–H) Relative expression of RAN reporters in HEK293 cells, HeLa cells, and rat primary hippocampal neurons after transfection with functionally capped (A**–**D) or nonfunctionally capped (E**–**H) mRNA for 24 hours. All data is normalized to FF and graphed relative to functionally capped AUG-NL reporter expression, n = 12. (I–N) Relative expression of C9 RAN reporters in HEK293 cells, HeLa cells, and rat primary hippocampal neurons after transfection with functionally capped (I**–**K) or nonfunctionally capped (L**–**N) mRNA for 24 hours. Data is expressed as NL signal normalized to FF and relative to GA70 reporter expression, n = 12. Graphs represent mean ± SEM; Brown-Forsythe one-way ANOVA with Dunnet’s multiple comparison correction. *P < 0.05, **P<0.01, ***P < 0.001, ****P < 0.0001. GA = poly-glycine-alanine, GP = poly-glycine-proline, GR = poly-glycine-arginine, +1CGG = poly-glycine, FF = firefly.

**Figure S2:**
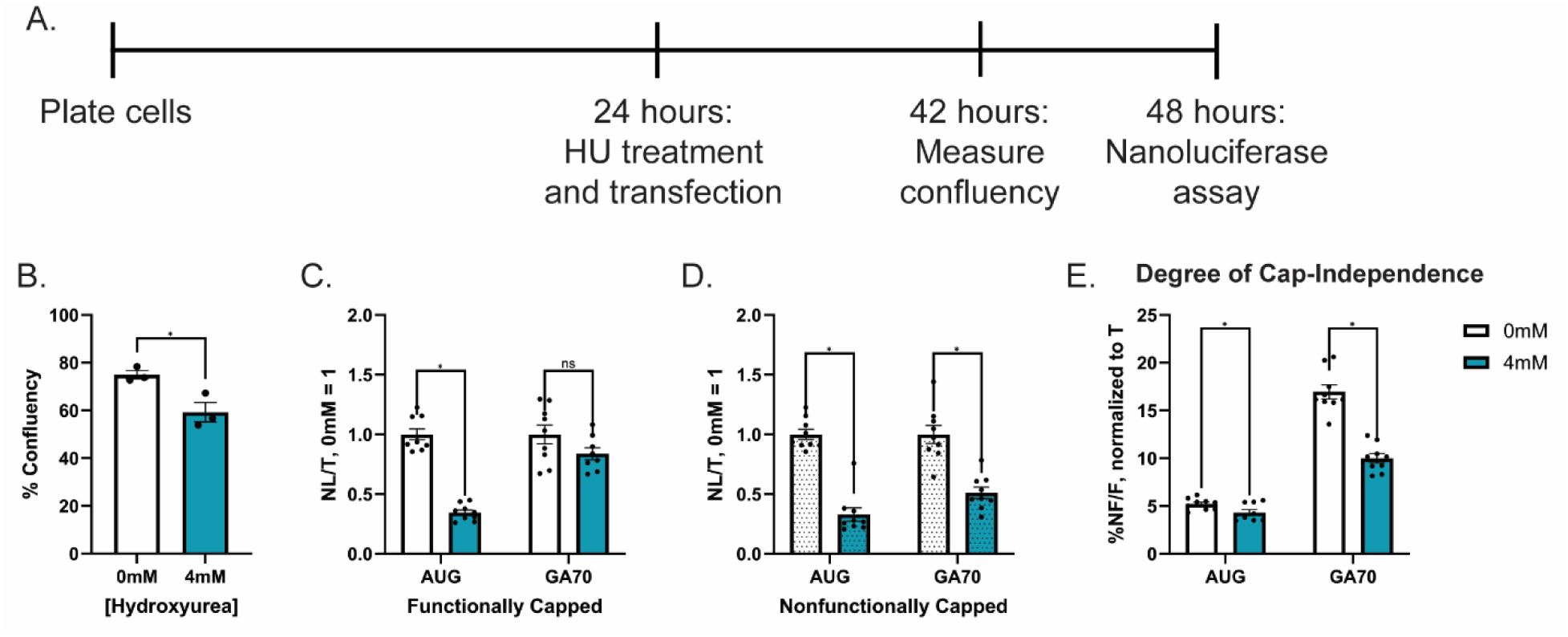
Cell division does not explain high cap-dependence of RAN translation. (A) Timeline of experiment. (B) Percent confluency of cells after treatment with hydroxyurea (HU) for 18 hours, n = 3 plates. (C-D) Relative expression of RAN reporters in HU or vehicle treated HEK293 cells after transfection with functionally capped (C) or nonfunctionally capped (D) mRNA for 24 hours. Data is expressed as NL signal normalized to titer, n = 9. (E) Degree of cap-independence of reporters after transfection into HU or vehicle treated HEK293 cells after transfection for 24 hours. Data is expressed as percent signal from nonfunctionally capped mRNA (NF) compared to the functionally capped mRNA (F), normalized to titer, n = 9. Graphs represent mean ± SEM; (B) Two-tailed Student’s t test with Welch’s correction. (C-E) Multiple Holm–Šídák’s two-tailed Student’s t test with Welch’s correction.*P < 0.05, **P<0.01, ***P < 0.001, ****P < 0.0001.

**Figure S3:**
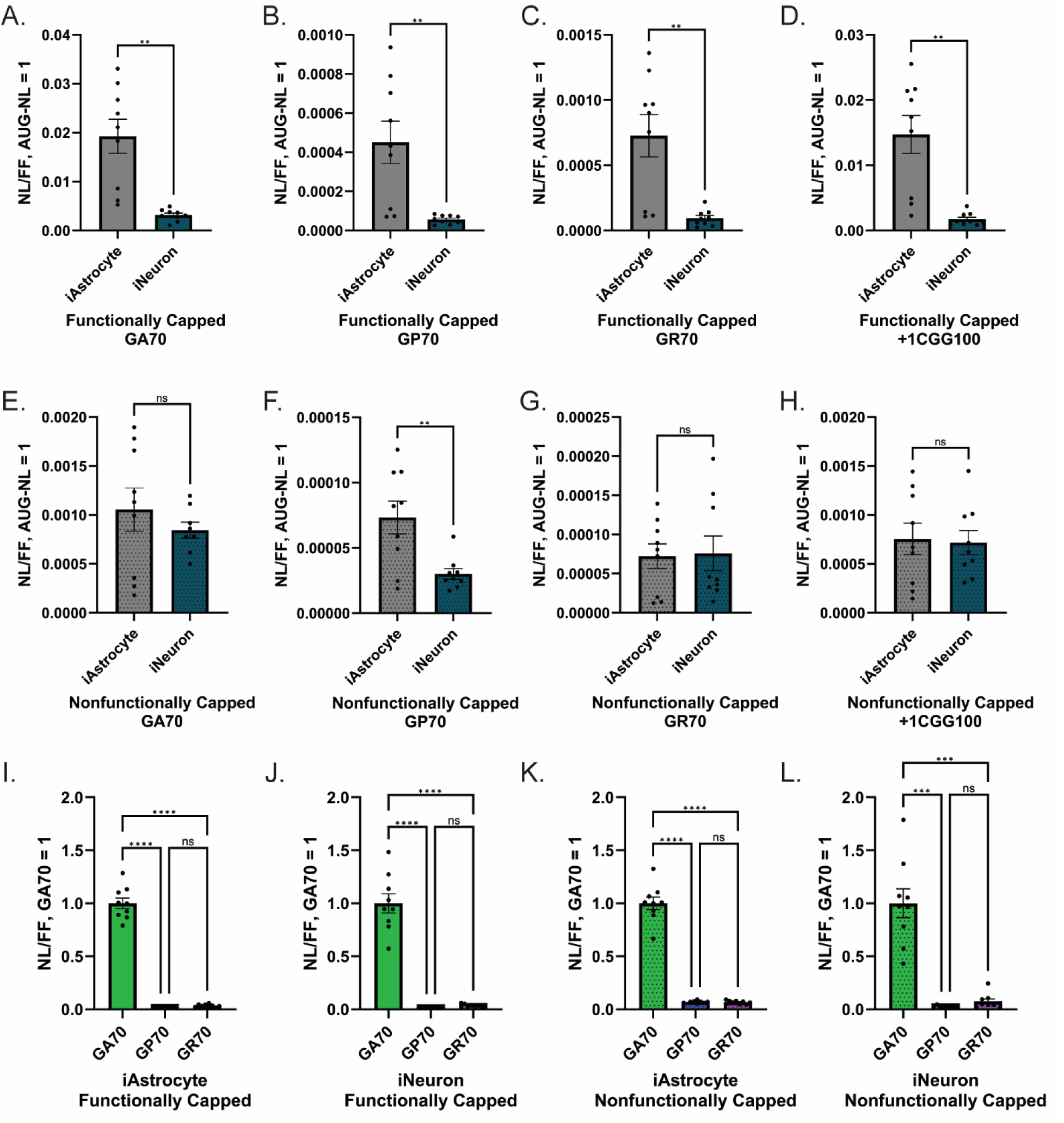
Expression of RAN reporters in human-derived cells. Relative expression of RAN reporters in iAstrocytes and iNeurons after transfection with functionally capped (A**–**D) or nonfunctionally capped (E**–**H) mRNA for 24 hours. Data is expressed as NL signal normalized to FF and relative to functionally capped AUG-NL reporter expression, n = 12. (I–L) Relative expression of C9 RAN reporters in iAstrocytes and iNeurons after transfection with functionally capped (I**–**J) or nonfunctionally capped (K**–**L) mRNA for 24 hours. Data is expressed as NL signal normalized to FF and relative to GA70 reporter expression, n = 12. Graphs represent mean ± SEM; (A**–**H) Two-tailed Student’s t test with Welch’s correction. (I**–**L) Brown-Forsythe one-way ANOVA with Dunnet’s multiple comparison correction. *P < 0.05, **P<0.01, ***P < 0.001, ****P < 0.0001.

**Figure S4:**
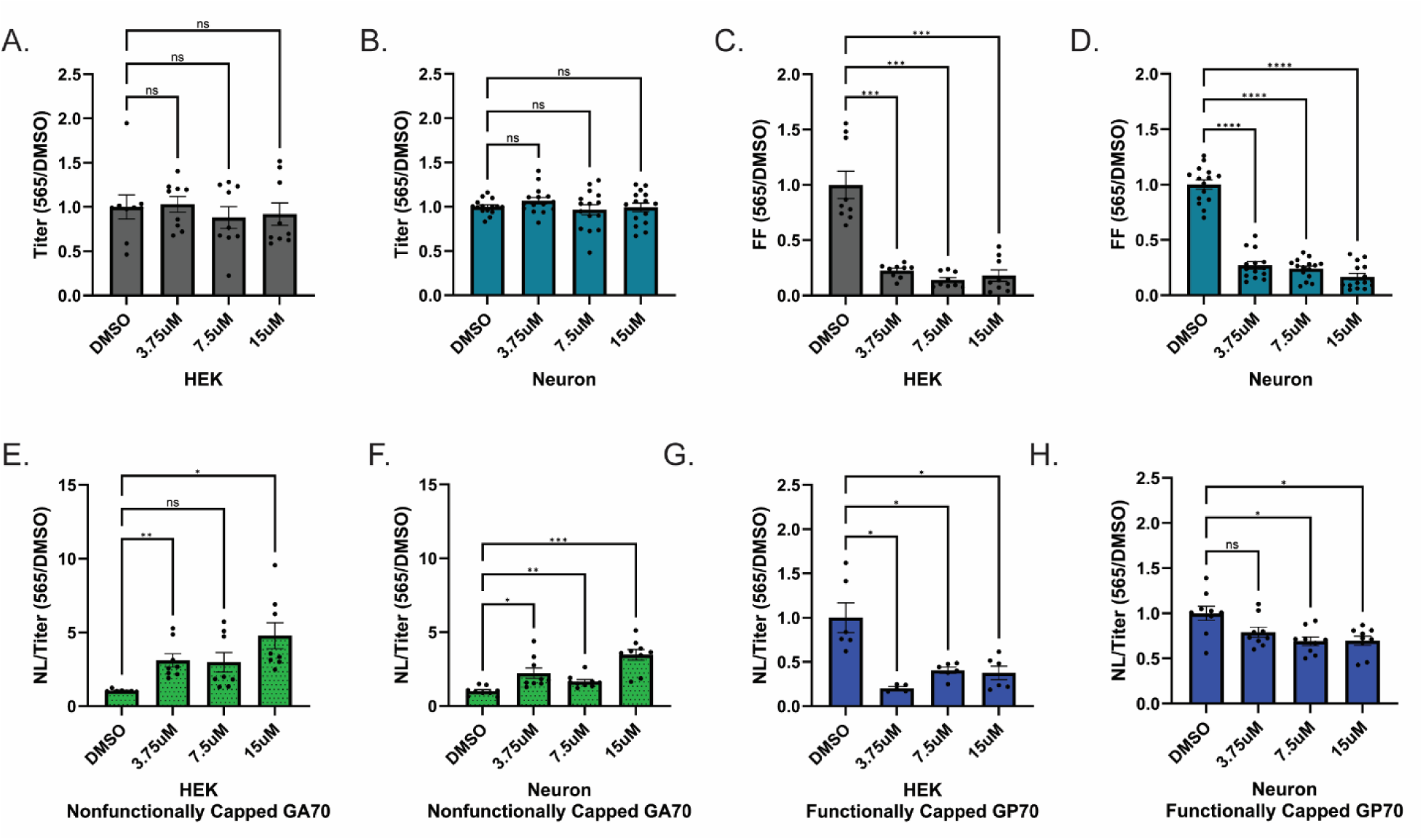
Effect of cap-binding factor eIF4E inhibition on non-canonical translation. (A–B) Cell viability measured by cell titer after 24 hours of treatment with 0 – 15μM eIF4E competitive inhibitor eFT-565 (565) in HEK293 cells (A) or primary rat hippocampal neurons (B) transfected with AUG-NL reporter 4.5 hours after treatment. Data expressed as titer signal relative to DMSO condition, n = 9 (A), n = 15 (B). (C–D) Effect on cotransfected firefly luciferase reporter signal after 24 hours of treatment with 0 – 15μM eFT-565 in HEK293 cells (C) or primary rat hippocampal neurons (D) transfected with AUG-NL reporter 4.5 hours after treatment. Data expressed as titer signal relative to DMSO condition, n = 9 (C), n = 15 (D). (E–F) Relative response of nonfunctionally capped GA70 to treatment with eFT-565. HEK293 cells (E) and rat primary hippocampal neurons (F) were treated with 0 – 15μM eFT-565, transfected with nonfunctionally capped GA70 reporter 4.5 hours after treatment, and assayed for translation 19.5 hours later. Data is expressed as NL signal normalized to titer and relative to DMSO treated condition, n = 9. (G–H) Relative response of GP70 to treatment with eFT-565. HEK293 cells (G) and rat primary hippocampal neurons (H) were treated with 0 – 15μM eFT-565, transfected with functionally capped GP70 reporter 4.5 hours after treatment, and assayed for translation 19.5 hours later. Data is expressed as NL signal normalized to titer and relative to DMSO treated condition, n = 6. Graphs represent mean ± SEM; Brown-Forsythe one-way ANOVA with Dunnet’s multiple comparison correction. *P < 0.05, **P<0.01, ***P < 0.001, ****P < 0.0001.

**Figure S5:**
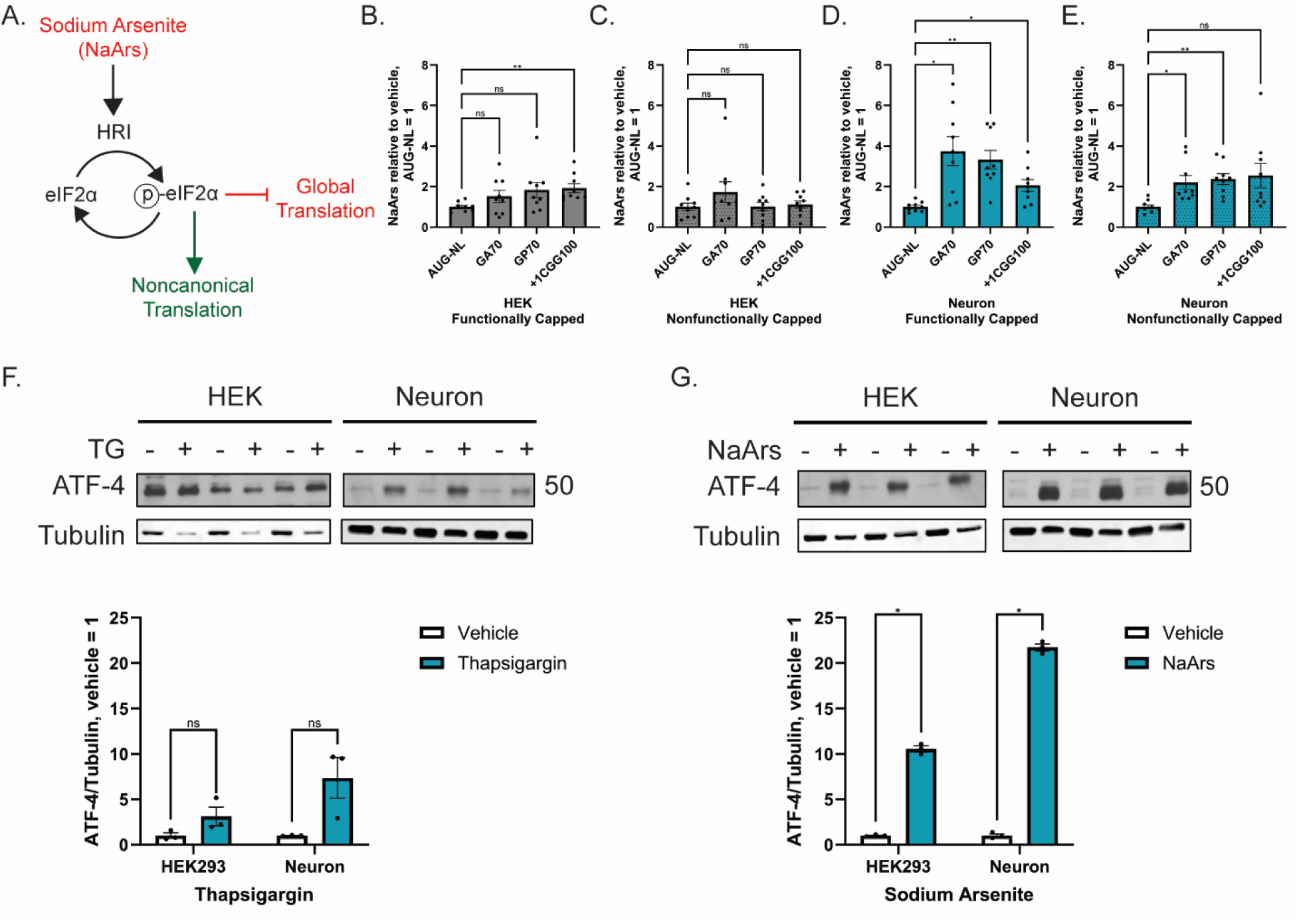
Cap-independent RAN translation is resistant to inhibition by sodium arsenite activation of the integrated stress response. (A) Schematic of the oxidative stress branch of the integrated stress response. (B–E) Relative expression of reporters in HEK293 cells (B-C) and rat primary hippocampal neurons (D**–**E) treated with 20μM sodium arsenite (NaArs), transfected with functionally capped (B/D) or nonfunctionally capped (C/E) reporters, and assayed for translation 5 hours later. Data is expressed as ratio of NL signal normalized to titer and relative to AUG response, n = 9. (F-G) Western blot and quantification of HEK293 cells and rat primary hippocampal neurons treated with 2μM thapsigargin (F) or 20μM NaArs (G) for 5 hours. Blots were probed for ATF-4 as a marker of the integrated stress response. Graphs represent mean ± SEM; (B-E) Brown-Forsythe one-way ANOVA with Dunnet’s multiple comparison correction. (F-G) Multiple Holm–Šídák’s two-tailed Student’s t test with Welch’s correction. *P < 0.05, **P<0.01, ***P < 0.001, ****P < 0.0001.

**Figure S6:**
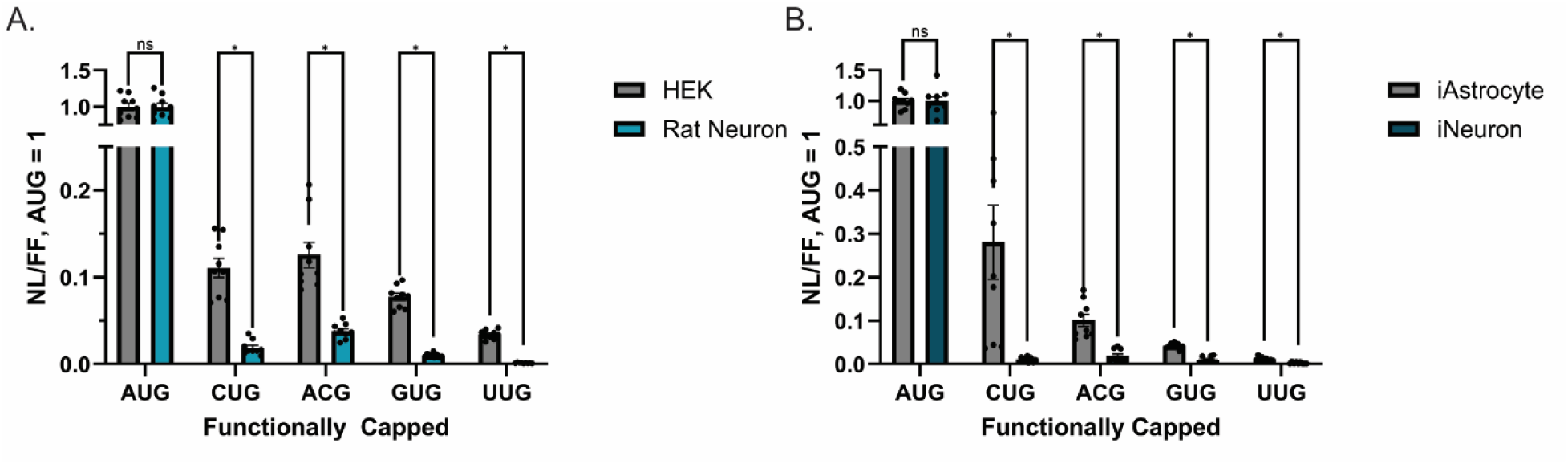
Start codon stringency across near cognate start codons is higher in neurons. Relative expression of near cognate codon reporters in HEK293 cells and rat primary hippocampal neurons (A) or iAstrocytes and iNeurons (B) after transfection with functionally capped mRNA for 24 hours. Data is expressed as NL signal normalized to FF with AUG-NL set to 1. Graphs represent mean ± SEM; Multiple Holm–Šídák’s two-tailed Student’s t test with Welch’s correction. *P < 0.05, **P<0.01, ***P < 0.001, ****P < 0.0001.

**Figure S7:**
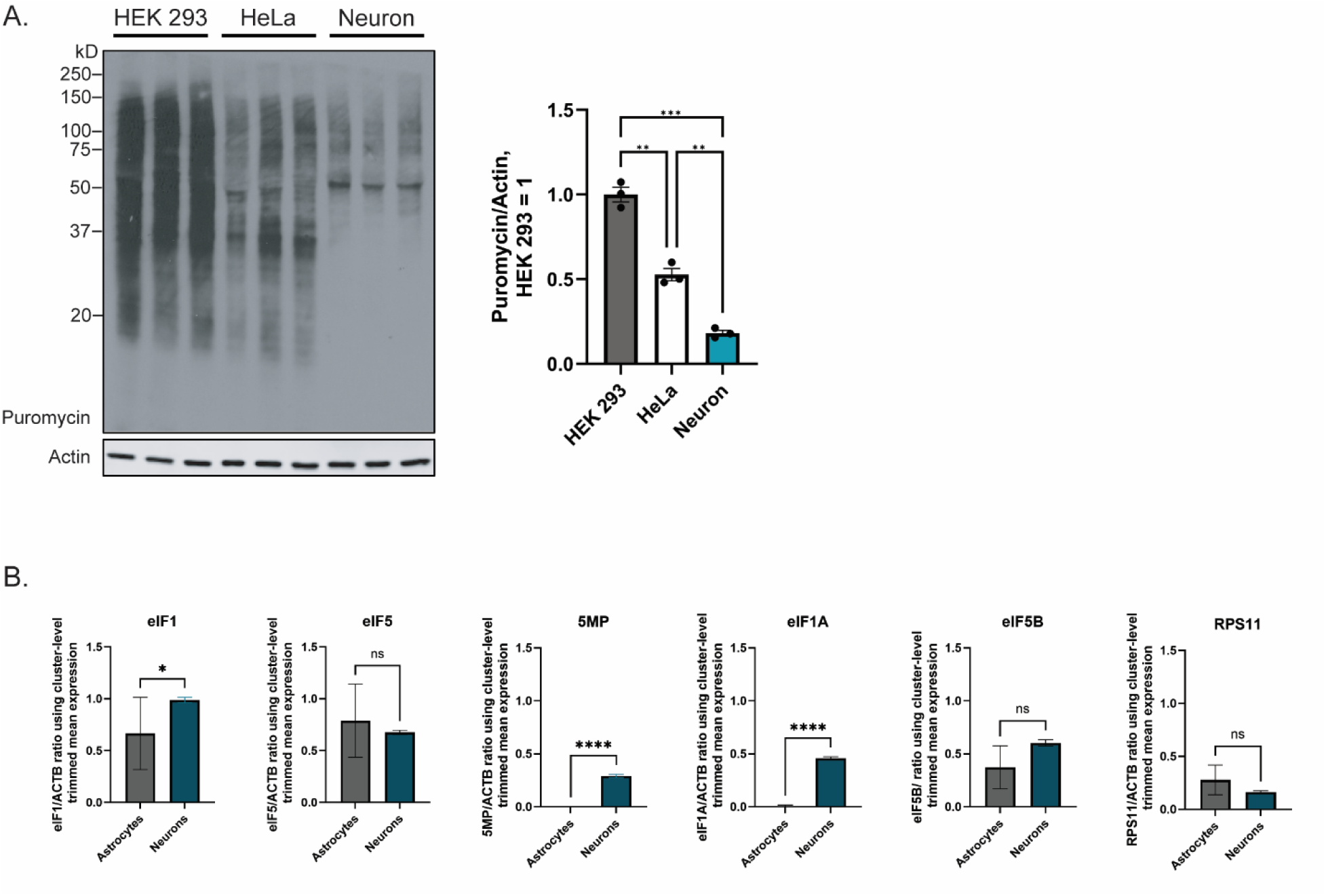
Initiation factor variation by cell type. (A) SUnSET assay and quantification of puromycin incorporation in HEK293 cells, HeLa cells and rat primary hippocampal neurons. Data is expressed as intensity in each condition normalized to actin (loading control), n = 3. (B) *In silico* quantification of initiation factor expression across cell types from scRNA seq GTEX datasets. The data utilized cluster-level trimmed means to compute ratios relative to ACTB. Transcriptomic clusters were grouped into Neurons, n = 226, and Astrocytes n = 6. Data is expressed as average trimmed-mean per group normalized to ACTB. Graphs represent mean ± SEM; (A) two-tailed Student’s t test with Welch’s correction; (B) Kolmogorov-Smirnov test. *P < 0.05, **P<0.01, ***P < 0.001, ****P < 0.0001.

**Table S1:**
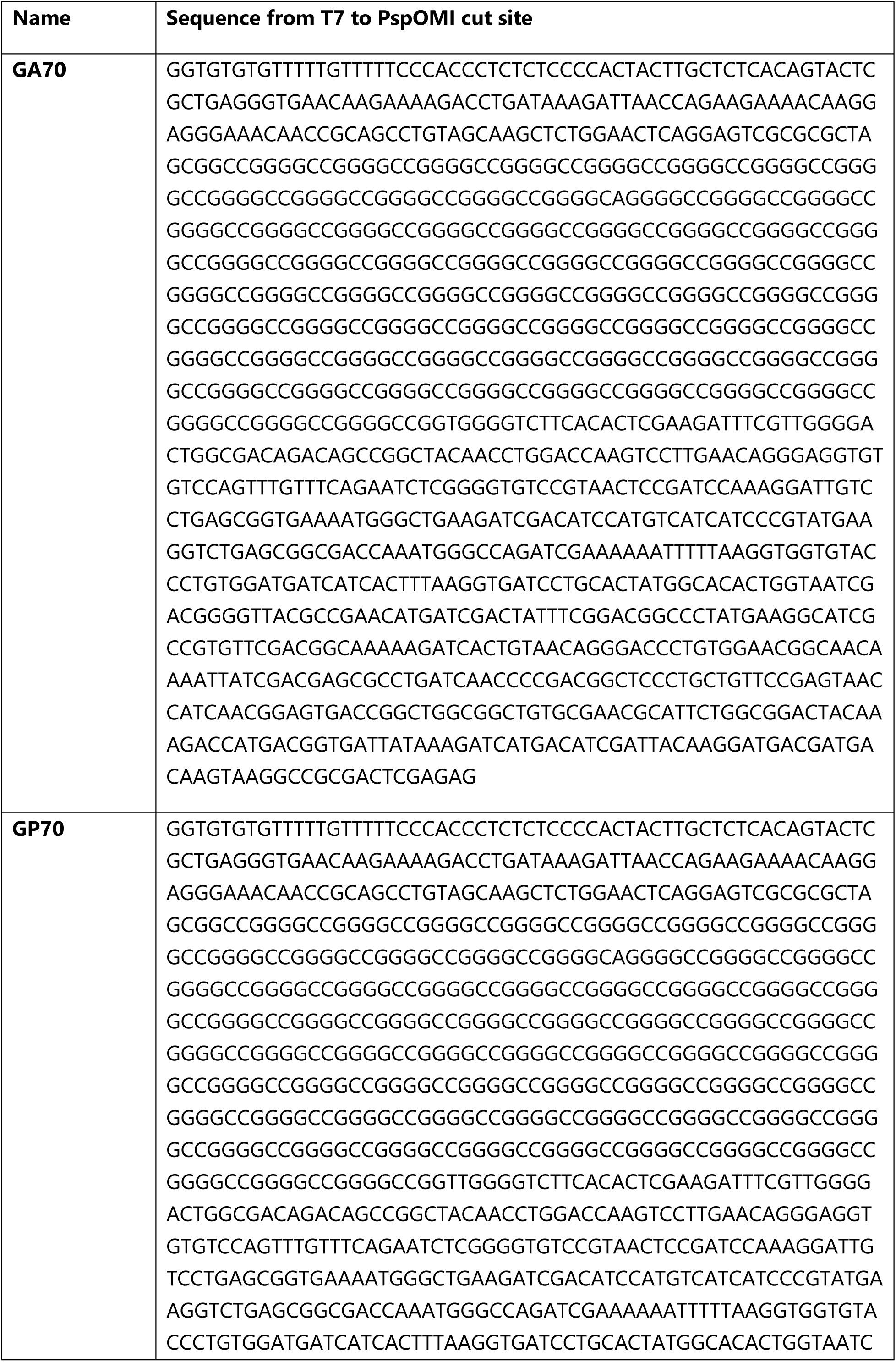
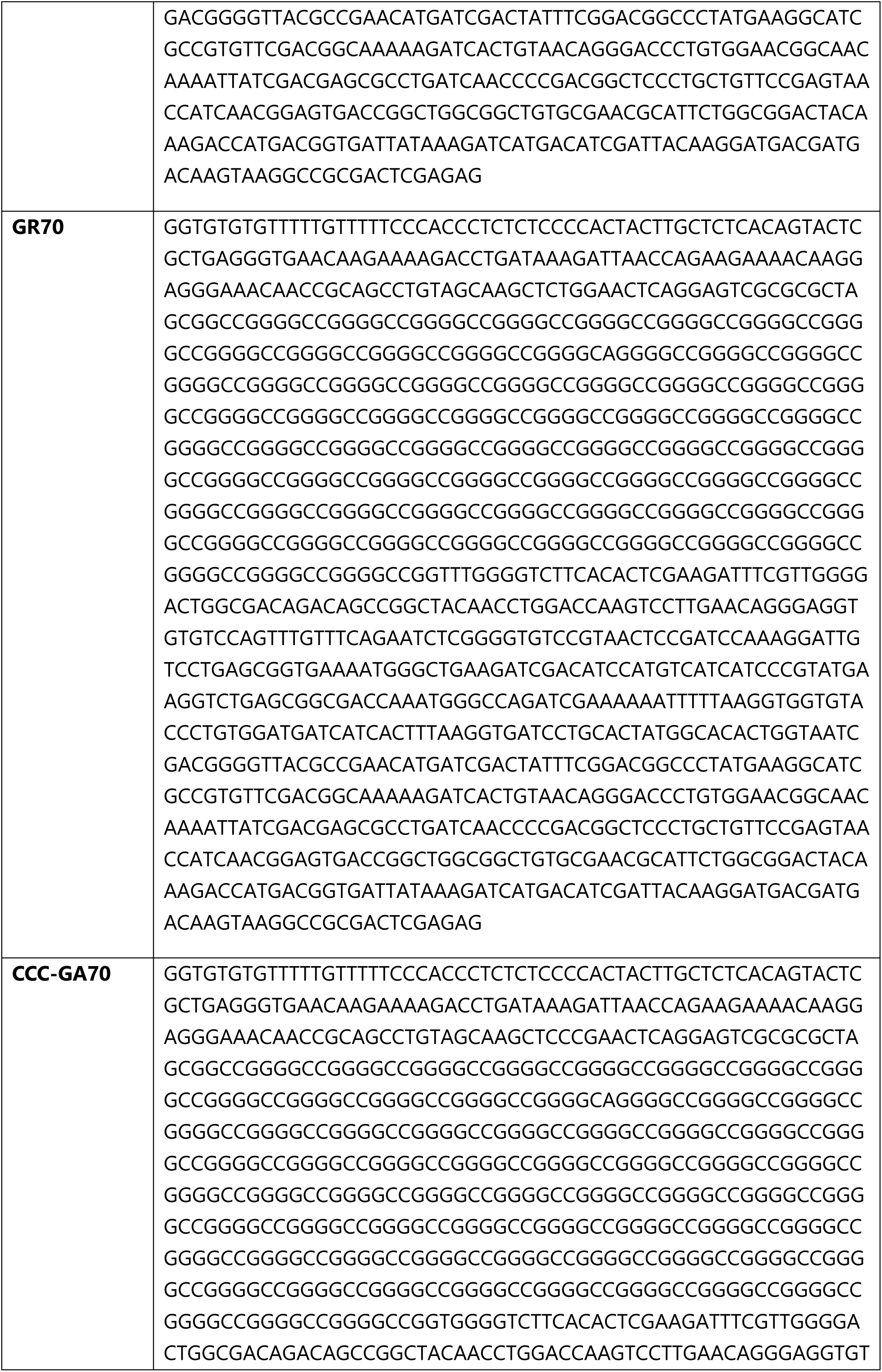
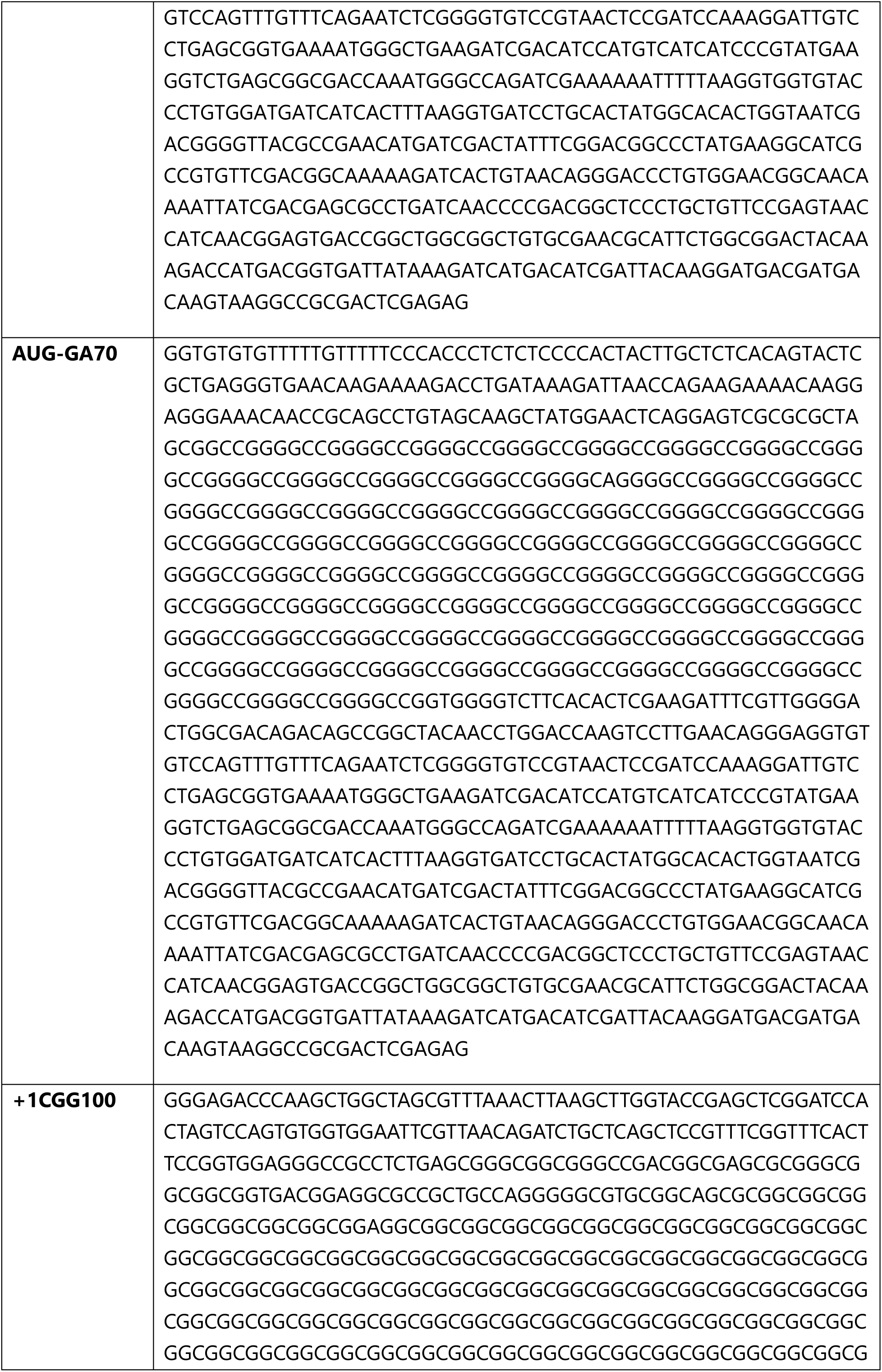
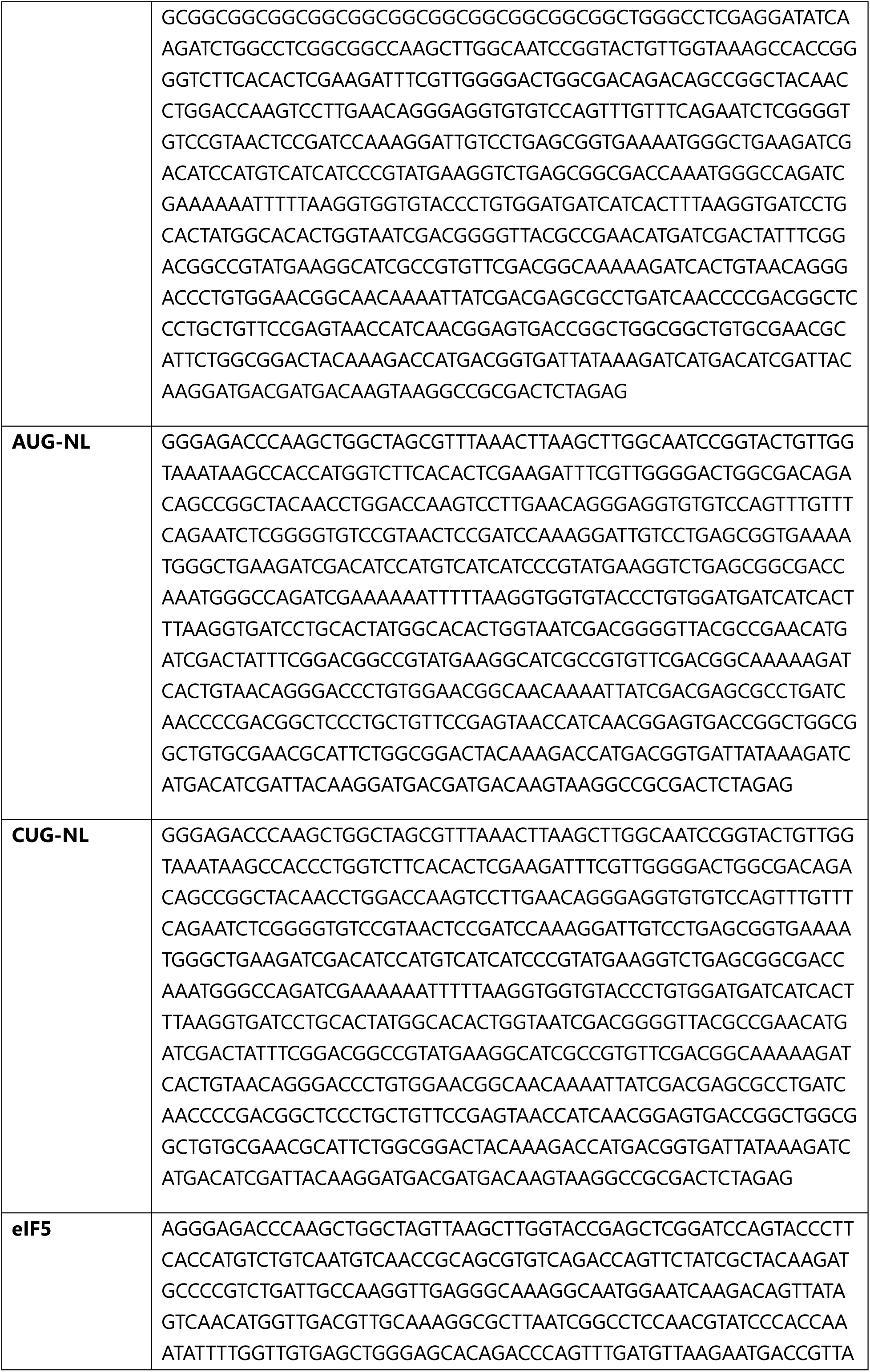
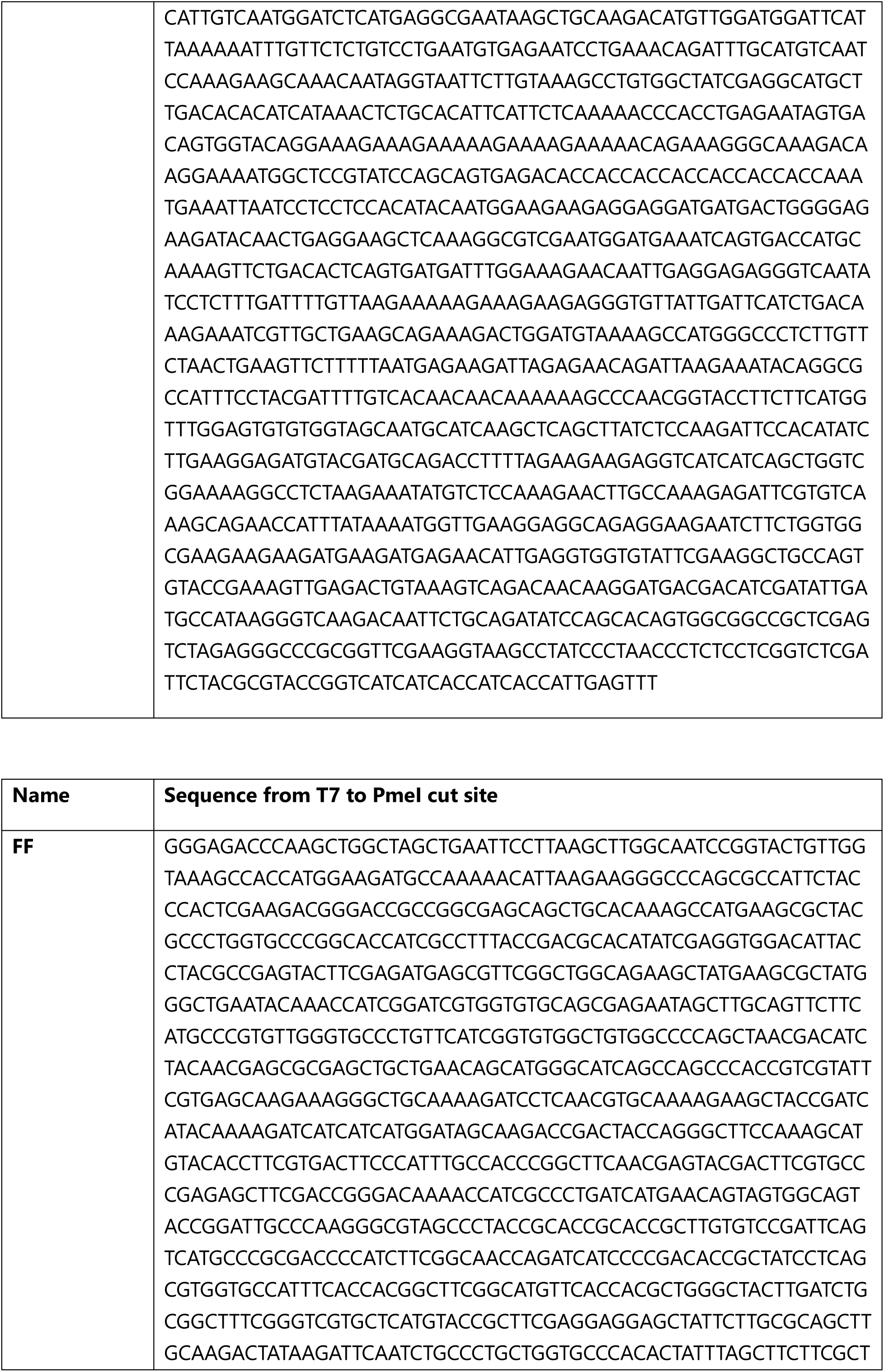
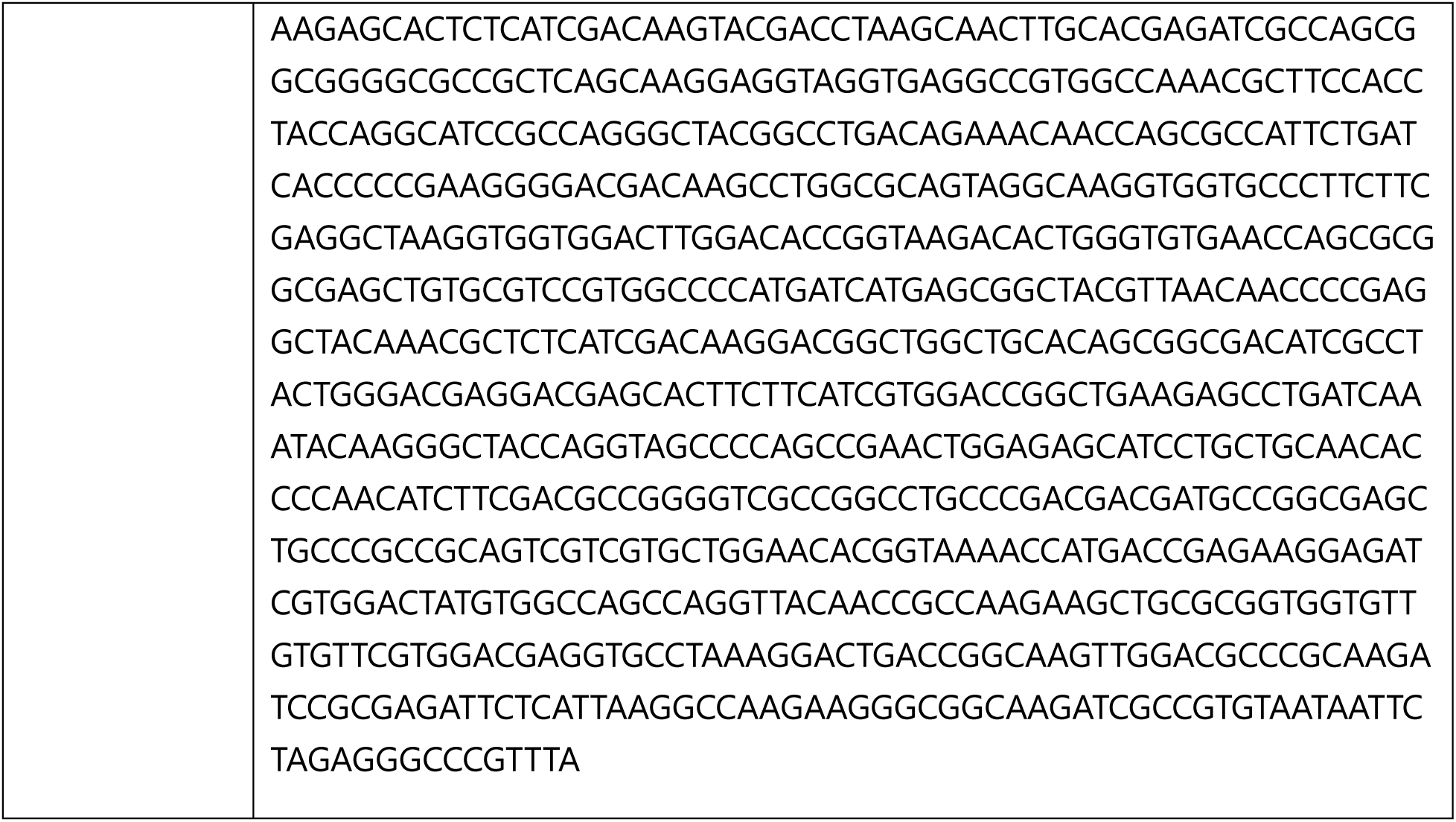
Reporter Sequences.

